# Multiplex single-molecule kinetics of nanopore-coupled polymerases

**DOI:** 10.1101/2020.03.15.993071

**Authors:** Mirkό Palla, Sukanya Punthambaker, P. Benjamin Stranges, Frederic Vigneault, Jeff Nivala, Daniel J. Wiegand, Aruna Ayer, Timothy Craig, Dmitriy Gremyachinskiy, Helen Franklin, Shaw Sun, James Pollard, Andrew Trans, Cleoma Arnold, Charles Schwab, Colin Mcgaw, Preethi Sarvabhowman, Dhruti Dalal, Eileen Thai, Evan Amato, Ilya Lederman, Meng Taing, Sara Kelley, Adam Qwan, Carl W. Fuller, Stefan Roever, George M. Church

**Author notes:** These authors contributed equally to this work (Mirkό Palla, Sukanya Punthambaker).

## Abstract

DNA polymerases have revolutionized the biotechnology field due to their ability to precisely replicate stored genetic information. Screening variants of these enzymes for unique properties gives the opportunity to identify polymerases with novel features. We have previously developed a single-molecule DNA sequencing platform by coupling a DNA polymerase to a α-hemolysin pore on a nanopore array. Here, we use this approach to demonstrate a single-molecule method that enables rapid screening of polymerase variants in a multiplex manner. In this approach, barcoded DNA strands are complexed with polymerase variants and serve as templates for nanopore sequencing. Nanopore sequencing of the barcoded DNA reveals both the barcode identity and kinetic properties of the polymerase variant associated with the cognate barcode, allowing for multiplexed investigation of many polymerase variants in parallel on a single nanopore array. Further, we develop a robust classification algorithm that discriminates kinetic characteristics of the different polymerase mutants. As a proof of concept, we demonstrate the utility of our approach by screening a library of ~100 polymerases to identify variants for potential applications of biotechnological interest. We anticipate our screening method to be broadly useful for applications that require polymerases with unique or altered physical properties.

## Introduction

DNA polymerases are enzymes that duplicate genetic information by synthesizing a new complementary DNA strand from the parent template, thereby preserving genetic information^1^. They are indispensable enzymes used in many biotechnological and clinical applications such as PCR, cloning, DNA sequencing, whole genome amplification and diagnostic testing^2,3^. There is a growing need for DNA polymerase variants with different efficacies, stabilities, processivities and fidelities, as well as for engineered polymerases with novel functions for specific uses^4^, for example, polymerases that can incorporate unnatural substrates^5–7^ or temperature sensitive mutants^8^. Screening a library of DNA polymerase mutants can provide novel candidates with unique properties for use in customized reactions^9^. To date, such mutants have been generated by directed evolution and methods for large-scale screening of polymerase variants using mutagenesis, phage display and compartmentalized self-replication^8,10–12^. This has led to the identification and development of different polymerases for a wide spectrum of applications in biotechnology^13,14^. However, these screening methods provide little to no information on the physical properties of each variant, for instance, their kinetic parameters. To obtain detailed kinetic characterization of polymerase mutants, these methods still rely on screening or selection of these enzymes one at a time (post-selection). Therefore, a multiplexed, single-molecule characterization method would be more beneficial.

Recently, single-molecule strategies have been shown to have great potential to monitor enzyme dynamics and to provide information about the molecular interactions that are hidden in ensemble measurements^15–20^. For example, single-molecule FRET studies have been used to determine protein interactions and enzyme kinetic parameters^21^. In Pacific Biosciences’ approach, a single DNA polymerase enzyme is immobilized at the bottom of a nanophotonic well, termed the zero-mode waveguide (ZMW), and single base incorporations are detected by fluorescence spectroscopy^22^. It has been shown that the fluorescence pulse width and interpulse duration could be used to monitor polymerase kinetics to detect DNA methylation^23^. Although ZMWs provide a high-throughput platform for single-molecule DNA sequencing, the signal has not been used to compare kinetics of different DNA polymerase mutants. Other classes of biosensing platforms are based on nanoscale field-effect-transistors (FET)^24–26^, which are designed for interrogating single molecules. While these sensor arrays may offer throughput enabled by wafer-scale fabrication, the yield of single-enzyme placement to a sensor element is currently limited to dilution-based approaches^18^. At best, this approach can only provide a Poisson distribution of populated sensors^27^. Additionally, the fabrication of these FET devices remains a manual and challenging process^18^.

We have previously shown that nanopore-based sequencing-by-synthesis (Nanopore-SBS) is a viable method for both sequencing and detection of single-molecule catalytic activity^28^. In that work, we had demonstrated the ability to insert individual α-hemolysin (αHL) nanopores into lipid bilayers on a complementary metal-oxide-semiconductor (CMOS) array, providing a highly-scalable platform for real-time and multiplex single-molecule enzyme measurements. The ionic current through the nanopore was measured using Ag/AgCl electrodes coupled to a silicon substrate integrated electrical circuit in voltage clamp mode. In our current study, we use a new CMOS chip containing thousands of individually addressable pores, which was developed by Roche Sequencing Solutions. Here, we have utilized polarizable platinum electrodes, where the measurement relies on non-faradaic conduction. We have also changed from using φ29 DNA polymerase to a homologue that has better performance and stability on the electrode array. Given these technological advances, here we present a method to monitor polymerase kinetics during single-molecule, real-time DNA sequencing. In this technique, single αHL-coupled polymerase molecules are observed while they catalyze the incorporation of tagged nucleotides complementary to a barcoded template DNA strand. The templates have circular topology, which enables multiple observations of the same barcoded region (**Fig. 1a**). Incorporation of a nucleotide is detected as a change in the voltage on the electrode when the tag specific for that nucleotide is repeatedly captured in the pore (**Fig. 1b**). Each tag generates a characteristic and well-separated signal, thus uniquely identifying the added base (**Fig. 1c**). The incorporation event ends when the tag is cleaved by the polymerase before moving to the next base in the DNA template. A variety of metrics related to tagged nucleotide incorporation and tag capture during the polymerase catalytic cycle can be measured in real-time, which adds valuable information about single-molecule DNA polymerase kinetics. Therefore, we hypothesized that kinetic parameters of polymerase variants might be distinguishable using our nanopore-based DNA barcode sequencing.

**Figure 1.**
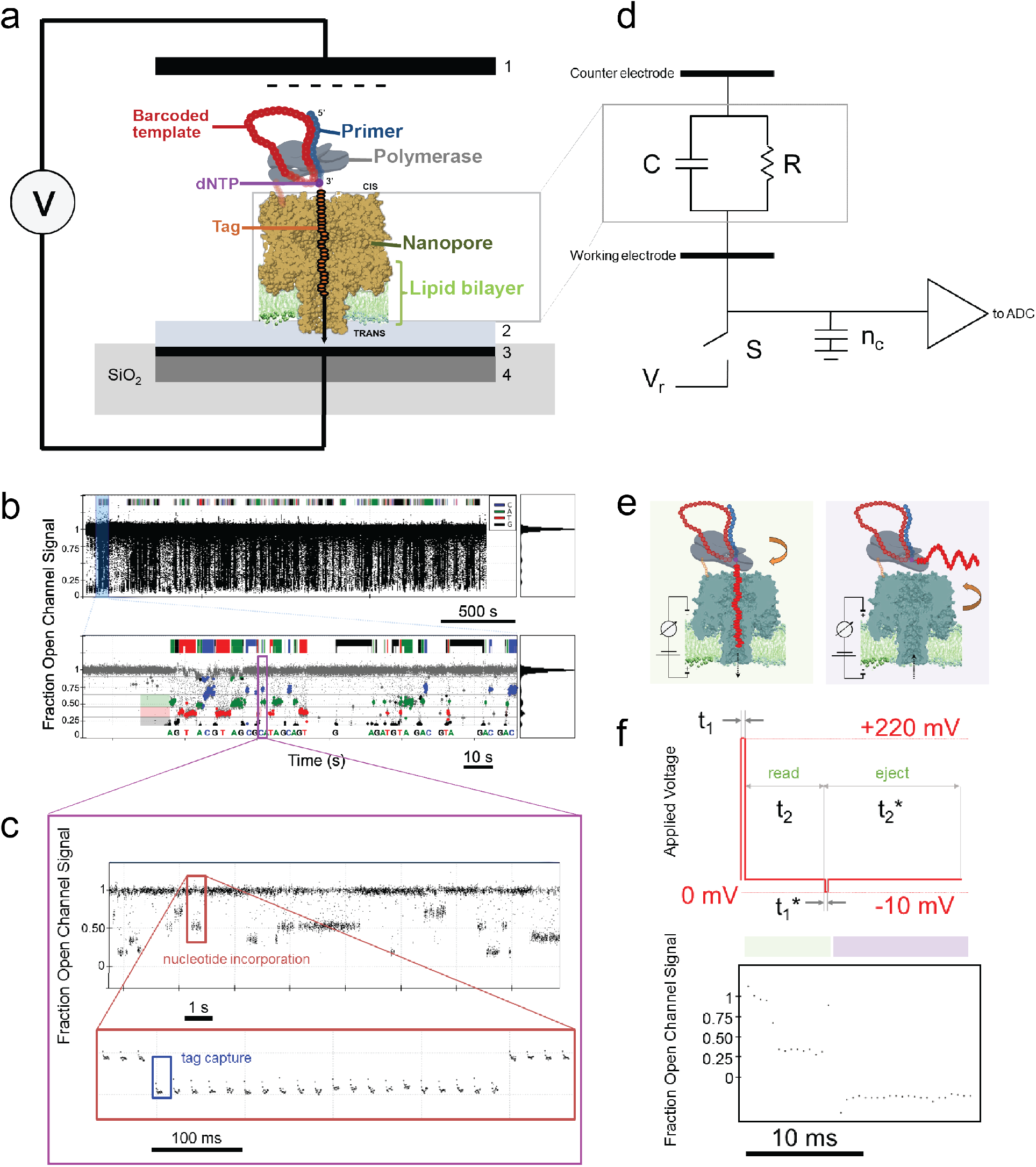
Principle of single-molecule circular barcoded DNA template sequencing on a nanopore array. (**a**) A DNA polymerase coupled to a αHL nanopore and loaded with a primed circular template is inserted into a lipid bilayer on a nanopore array. Sequencing starts by adding tagged nucleotides that provide a characteristic voltage signature during incorporation as the tag is repeatedly captured in the pore. Measurement setup contains a counter electrode (**1**), an analog measurement circuitry (**4**) connected to a platinum working electrode (**3**), which is covered by a thin film of electrolyte solution (**2**). (**b**) Fraction open channel signal (FOCS) *versus* time trace of tagged nucleotide captures for a single pore during a typical DNA sequencing experiment [top panel]. The identified base calls are highlighted in standard Sanger colors in a zoomed-in region [bottom panel]. A histogram of FOCS is shown in the right sub-panels. The dominant peak corresponds to the open channel signal (FOCS=1), while the four minor peaks represent signal associated with each tagged nucleotide capture events. (**c**) Tag captures [blue box, bottom panel] are detected by measuring the conductance of the pore during a single nucleotide incorporation [red box, top panel]. Data points collected only during the positive voltage commands (“read” periods) are displayed. (**d**) Schematic representation of RC equivalent circuit of a pore-polymerase complex inserted in the lipid bilayer. In this circuit (grey box), the capacitor (C) represents the membrane capacitance, while the resistor (R) represents the resistance associated with the nanopore. (**e**) Schematics of a tag capture/ejection. The left panel represents a “read” period when a tag is being captured in the pore. The right panel represents an “eject” period when the same tag is being ejected from the pore. (**f**) Dynamic voltage control. A brief signal pulse (+220 mV for t_1_) is applied across the membrane, which captures the tag in the pore and charges the membrane capacitance. Then, the membrane is discharged, defined as “read” period (0 mV for t_2_), during which the signal decay is recorded with a sampling rate of 2 kHz. The voltage is then reversed (−10 mV for t_1_*), which ejects the tag from the pore. Next, during the “eject” period (0 mV for t_2_*) the conductance decay is sampled as before [top panel]. The corresponding FOCS response is shown below the control signal [bottom panel].

## Results

### Principle of electrical recording

Sequencing experiments were performed using a CMOS chip that contained individually addressable platinum electrodes coupled to a silicon substrate integrated electrical circuit (**Fig. 1d**). These electrodes allowed voltages to be applied only to the membranes in specific wells, thus permitting independent sequence reads at these locations. The zoomed-in region (**Fig. 1d**, grey box) is a schematic representation of RC equivalent circuit of a pore-polymerase complex inserted in the lipid bilayer. Here, the capacitor (C=100 fF) represents the membrane capacitance, while the resistor (R) represents the resistance associated with the nanopore. A voltage source (V_r_) was selectively connected to or disconnected from C using a switch (S) controlled by a reset signal. When the voltage source was connected, C was charged. When the voltage source was disconnected, C was discharged through the nanopore. The voltage decay (τ=RC) was recorded with a sampling rate of 2 kHz, which was used to identify the different states of the nanopore. The unblocked state (**Fig. 1e**, right panel) corresponds to the open channel reading when no tag was captured in the pore and the blocked state (**Fig. 1e**, left panel) corresponds to having a tag captured in the pore (R=~10 GΩ). At each sampling point, the voltage across the second capacitor (n_c_=40 fF), that was placed parallel with the RC circuit, was measured by an analog-to-digital converter (ADC). Noise in the signal was low-pass filtered at 200 Hz cutoff frequency. The recorded ADC values were normalized (**Methods**) and were subsequently reported as fraction open channel signal (FOCS).

### Dynamic voltage control

A dynamic voltage command was used to interrogate the tag captures during a nucleotide incorporation, which provided a non-faradaic AC modulation of a rectangular wave (V_max_ = +220mV, V_min_ = −10 mV) with a 40% duty cycle and a frequency of 50 Hz applied across the lipid bilayer (**Methods**). The dynamic voltage control had two distinct stages: (1) a “charge” period, when a short voltage pulse was applied to charge the membrane capacitance and (2) a “discharge” period, when the membrane was discharged, during which the applied voltage was zero. Therefore, there was a brief signal pulse applied (with duration of t_1_) followed by an ADC response (with duration of t_2_) (**Fig. 1f**, top panel) during which the signal decay was recorded with a sampling rate of 2 kHz (**Fig. 1f**, bottom panel). First, a short positive voltage pulse (+220 mV for t_1_) was applied to charge the membrane capacitance and it was immediately discharged. The voltage decay recorded during this time (0 mV for t_2_) was defined as the “read” period. Next, a voltage pulse with negative polarity (−10 mV for t_1_*) was applied, which was followed by the “eject” period (0 mV for t_2_*) (**Fig. 1f**). During a sequencing experiment, the “read” and “eject” commands were continuously alternated to repeatedly interrogate the same tagged nucleotide during incorporation as well as to determine open channel conditions inferring the absence of incorporation activity.

### Sequencing of unique templates

To test if we could identify circular templates using the polymerase-nanopore system, we designed three synthetic single-stranded DNA (ssDNA) molecules consisting of a unique 32-nt barcode region flanked by a common 19-nt primer region (**Supplementary Fig. 1a**). They were circularized using either CircLigase or T4 ligase utilizing the primer region as a splint, then primed with the same universal primer to generate the circular barcoded templates (CBT) (**Supplementary Fig. 1b,c**). All CBTs met two design specifications, (1) all sequence identities were <85% when the templates were locally aligned to each other to make them serve as unique identifiers, and (2) the structures were optimized to eliminate regions of high base-pairing probability after circularization (**Supplementary Fig. 1d-f**). We used three different ϕCPV4 DNA polymerase variants (henceforth referred to as RPol), with mutations, identified during directed-evolution experiments (**Methods**). Pore-polymerase conjugates were complexed with each of the three unique circularized DNA templates (RPol:CBT) (**Methods**), which were finally loaded onto the chip for nine separate sequencing runs.

To measure the change in voltage through the nanopore, we employed a CMOS chip containing 32,768 individually addressable electrodes (**Methods**). Measurements were sampled at a rate of 2 kHz with a 40% duty cycle and AC frequency of 50 Hz by applying an alternating square waveform (+220 mV/−10 mV) across the lipid bilayer (**Methods**), which enabled the repeated interrogation of the same, tagged nucleotide during incorporation. Sequential nucleotide additions were detected as continuous tag captures associated with each of the four tagged nucleotides at characteristic signal levels through the pore (**Supplementary Fig. 2a**). Each tag generated a distinct and well-separated signal, uniquely identifying the added base (**Supplementary Fig. 2b**). The recorded signal levels were converted to raw reads using a probabilistic base caller software (**Methods**) after data acquisition in offline mode. We collected over 1,000 high-quality raw reads (**Methods**) for each RPol:CBT combinations (**Supplementary Table 1**) and observed multiple full iterations around the circular templates. These results confirmed that we could load polymerases with circular templates and sequence these templates. This showed the feasibility of template identification on the CMOS chip.

### Barcode identification

To demonstrate the suitability of barcode identification, we implemented a Smith-Waterman alignment-based barcode classification algorithm (**Methods**), which computes a probability score, henceforth defined as barcode match probability index (BMPI), that describes a relative measure of how uniquely a barcode can be identified compared to the other possible barcodes in the measurement set. First, high-quality reads were filtered out by requiring their read length to be greater than one (51 bp) and less than ten full barcode iterations and their consensus sequence length to be greater than 10 bp (**Methods** and **Supplementary Fig. 3**). Then, we used this classifier to analyze the RPol1:CBT1 sequencing data for estimating the accuracy with which one could identify the loaded barcoded DNA template. When the filtered raw reads were compared to the correct template (CBT1), the mean of the calculated BMPI values was 0.85 (**Fig. 2a**, left panel). In contrast, when the same reads were aligned to the incorrect templates (CBT2 and CBT3), their average BMPI values decreased to ~0.65 (**Fig. 2a**, left panel). Using this barcode identification strategy, a similar classification was performed by analyzing the RPol1:CBT2 and RPol1:CBT3 sequencing datasets, respectively. For both cases, the mean BMPI value was >0.80 when the raw reads were compared to the correct template and <0.80 when compared to the incorrect ones (**Fig. 2a**, middle and right panels). Similarly, as shown for CBT1, both CBT2 and CBT3 uniquely identified the polymerase variant based on the sequencing alignment metrics established above. Next, sequencing datasets for the other two pore-polymerase variants (RPol2, RPol3), each loaded with the three unique circular DNA templates, were similarly classified as for RPol1 described above. For all cases, we successfully identified the barcoded templates loaded on the polymerase variants (**Fig. 2b-c**). To further test the viability of our classifier, by computing a confusion matrix (**Supplementary Table 2**), we determined that when the BMPI value was >0.80 for a particular raw read, there was only ~2% probability of misidentifying the barcode. For this reason, we chose 0.80 BMPI as a threshold value to identify barcodes with high confidence. These findings demonstrate that with reads >50 bases, the BMPI value enables us to identify what template is bound to the polymerase.

**Figure 2.**
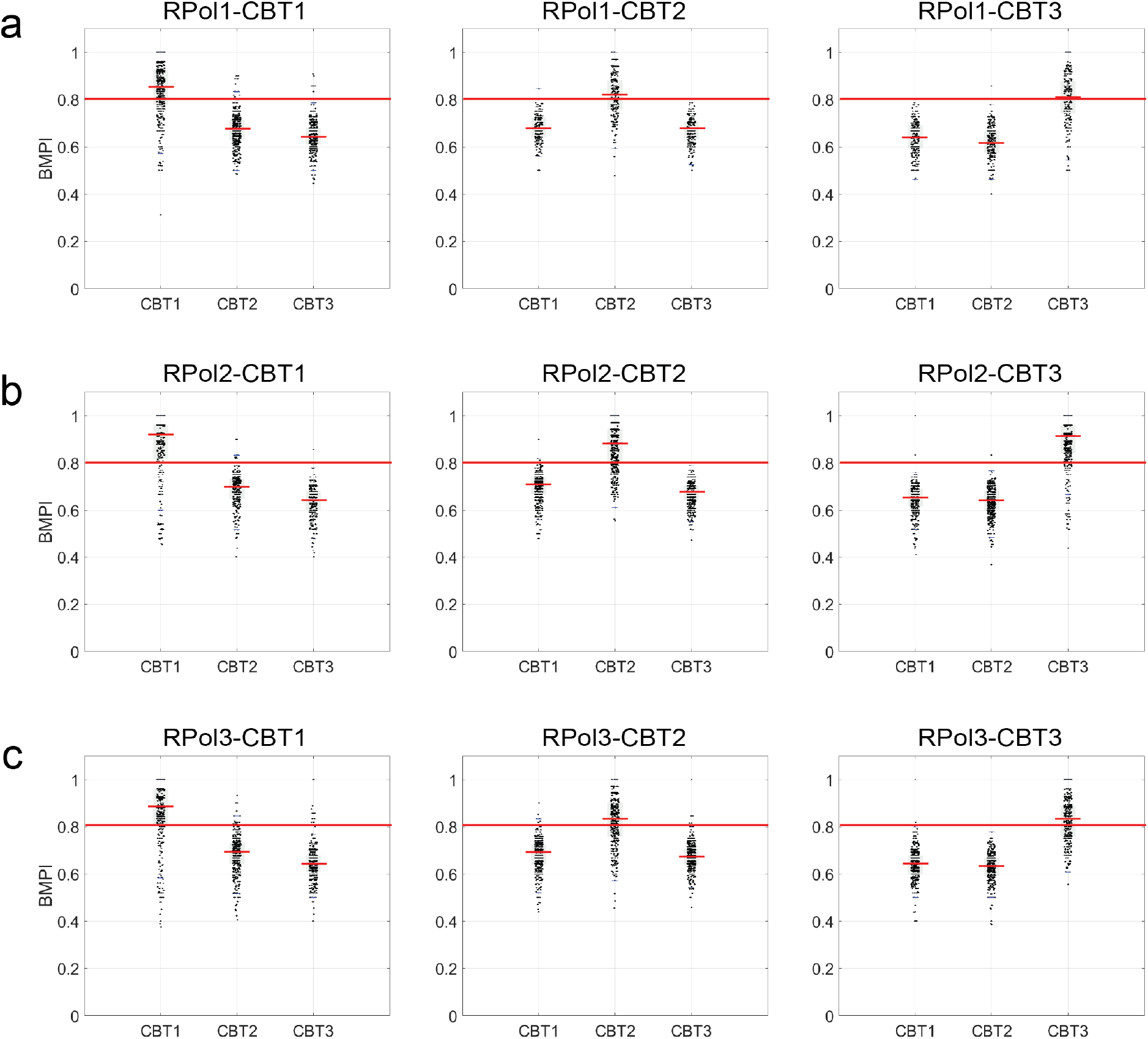
Barcode identification on a nanopore array. Barcode match probability index (BMPI) values of the three polymerase variants (**a**: RPol1, **b**: RPol2 and **c**: RPol3) loaded with the three unique DNA templates (CBT1, CBT2 and CBT3) calculated by the alignment-based barcode classifier (**Methods**). In each panel, barcode classification is shown when the high-quality raw reads are aligned to the correct and incorrect barcodes. For every RPol:CBT combination, the mean BMPI value was >0.80 when the raw reads were compared to the correct template and <0.80 when compared to the incorrect ones. A red line denotes the 0.80 BMPI cutoff. In each distribution, the red central mark indicates the mean. Raw data are jittered along the x-axis for clarity.

### Polymerase-barcode assignment is unique

After confirming that DNA templates loaded on each polymerase can be identified on the CMOS chip, we sought to determine if a template could be replaced with a different template once the pore-polymerase-template complex was formed. This was important because template replacement would make our barcoding strategy fail. To verify that replacement does not occur, we assembled the RPol2:CBT2 complex, which was subsequently loaded onto the chip for four different sequencing runs. First, we carried out a control run, in which the tagged nucleotides were added only after pore insertion. By employing our barcode classifier, we found that when the raw sequencing reads were compared to the correct template (CBT2), the mean BMPI value was 0.85 (**Supplementary Fig. 4**, Experiment 1). In contrast, when the same reads were aligned to an incorrect template (CBT1), this value decreased to ~0.70 (**Supplementary Fig. 4**, Experiment 2). As shown before, this confirmed that 0.80 BMPI can be used as a threshold value for barcode identification. Next, in the second set of experiments, we spiked in a 5-fold molar excess of a secondary barcode (CBT1) immediately after the pore-polymerase-template assembly, which mimics a multiplex scenario with a set of barcodes present in the same reaction volume during assembly. In two separate experiments, this complex was inserted into the membrane after a brief (<5 min) and after an overnight (~12 hr) incubation period, which provided two different time intervals for the added secondary template (CBT1) to replace the primary template (CBT2) already bound to the pore-polymerase complex. Then, tagged nucleotides were added to the subsequent sequencing reaction. For both cases, the mean BMPI value was >0.80 when the raw reads were compared to the template originally linked to the complex (CBT2) (**Supplementary Fig. 4**, Experiments 3 and 5) and <0.80 when compared to the template spiked in later (CBT1) (**Supplementary Fig. 4**, Experiments 4 and 6). Our results demonstrated that, even, after an overnight incubation with a second barcode, no barcode replacement took place. Additionally, we tested the possibility of on-chip barcode replacement, which mimicked a scenario with multiple barcodes present in the same reaction volume in the *cis* chamber of the CMOS chip. To this aim, when we spiked in a second barcode (CBT1) along with the tagged nucleotides after pore insertion, our barcode classification results indicated that our polymerase variants are uniquely labeled with their respective barcodes. Again, the mean BMPI score was above (**Supplementary Fig. 4**, Experiment 7) and below (**Supplementary Fig. 4**, Experiment 8) the threshold value of 0.80 for the primary template (CBT2) and the secondary template (CBT1), respectively. This confirmed that once a polymerase is loaded with a barcoded template it is not replaced by another template, *i.e.*, the polymerase-barcode assignment is unique.

### Kinetic properties of polymerases

One might be interested to screen for a particular DNA polymerase mutant having a defined set of kinetic properties characterized by enzyme fidelity, processivity, elongation rate, or lifetime. Multiplexed screening for these properties in parallel at the single-molecule level is not possible with current methods. In this system, a variety of kinetic parameters related to tagged nucleotide incorporation and tag captures can be derived from the voltage signal produced by single-molecule events during the polymerase catalytic cycle. For each base, we define dwell time (t_dwell_) as the duration of continuous “read” periods that found the pore blocked (**Fig. 3a**, cyan double arrow). Additionally, we define full catalytic rate (FCR) as the frequency of the full catalytic cycle (**Fig. 3a**, purple double arrow), which is the total time from the first “read” period that finds the pore blocked until the next one that finds it unblocked again. These parameters add valuable information about DNA polymerase kinetics as t_dwell_ is correlated with the total time required for a distinct nucleotide to be incorporated into the template, and FCR is determined by the kinetics of two successive catalytic events, tagged nucleotide incorporation and tag cleavage by the polymerase^28^. As an initial test, we calculated these kinetic parameters for each of the three polymerase variants loaded with a unique CBT from the already collected sequencing data shown in **Fig. 2**. When comparing the three different polymerase mutants, each loaded with the same template, we found that the mean FCR was ~0.7 s^−1^ for RPol1, ~1.5 s^−1^ for RPol2 and ~2.1 s^−1^ for RPol3 for all of the four bases (A, C, T, and G) regardless of the sequence context of the barcoded DNA template (**Fig. 3b** and **Supplementary Table 3**). Similarly, analysis of the mean dwell time of the tagged nucleotide incorporations were also independent of barcode content with computed values of ~1.2 s for RPol1, ~0.7 s for RPol2 and ~0.5 s for RPol3, respectively (**Supplementary Fig. 5**). These results demonstrated that the kinetic parameters are statistically different for each of the polymerase variants and that they are independent of barcode sequence context (**Fig. 3b**). For this reason, sequencing data for each of the three polymerase variants loaded with different templates were lumped into the same dataset for downstream analysis. This allowed us to classify polymerase kinetics based on template identification.

**Figure 3.**
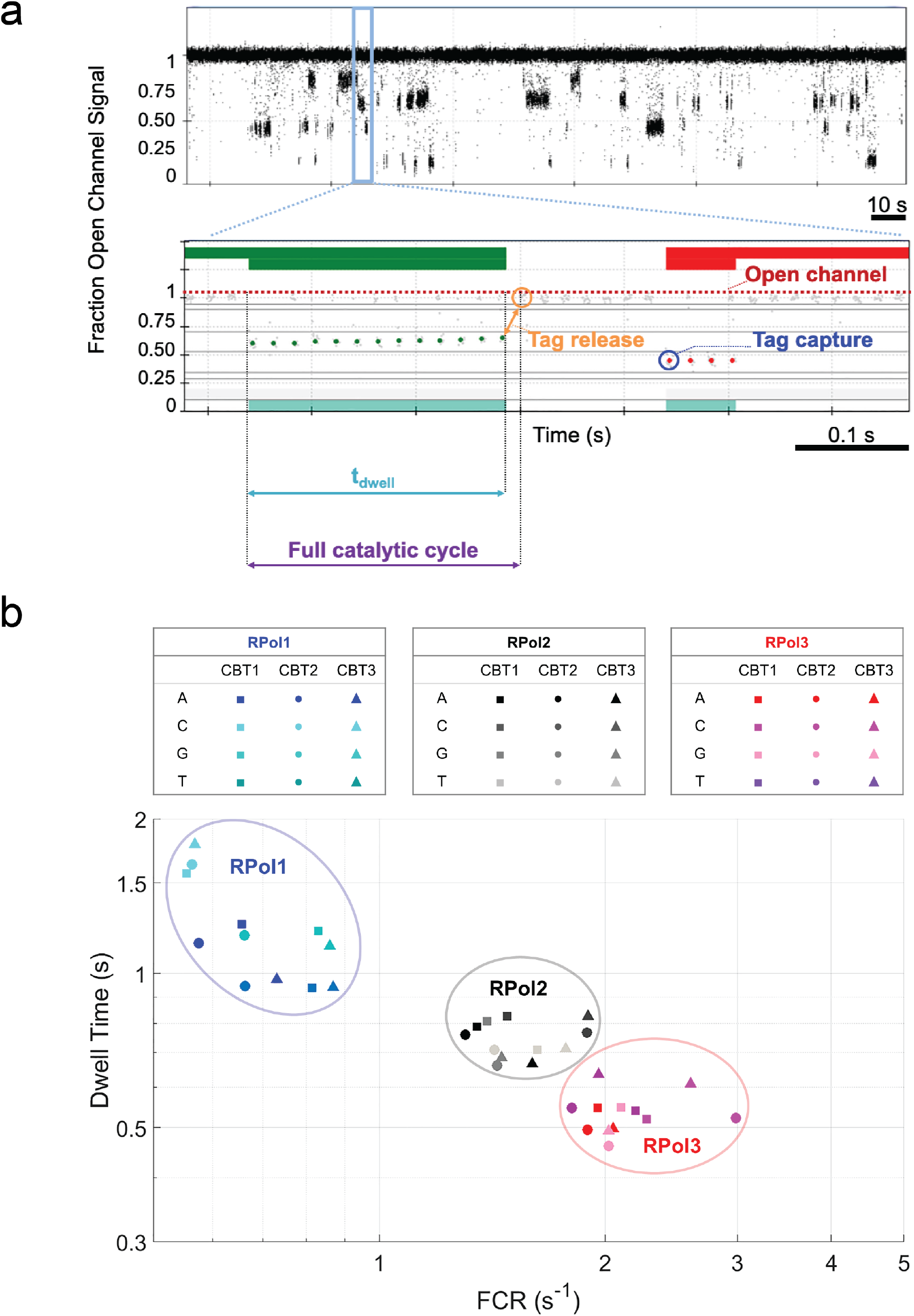
Kinetic properties of polymerase variants. (**a**) Definitions of key kinetic properties derived from single-molecule tagged nucleotide capture signal. Tagged nucleotide incorporation events detected by the nanopore [top panel]. Two incorporation events are highlighted in standard Sanger colors in a zoomed-in region [bottom panel, green and red blocks]. Within these blocks, the green and red points represent the tag capture events during a distinct nucleotide incorporation. Open channel signal (FOCS=1) is indicated in red dotted line. The first tag capture is indicated by the green circle at FOCS=0.6 followed by repeated interrogations of the same tag, followed by multiple tag capture events (stream of green points) and the tag release (orange double arrow). (**b**) Bird’s-eye view of the polymerase variant kinetics. Each marker represents the mean full catalytic rate (FCR) and mean dwell time (t_dwell_) value pair corresponding to each of the RPol:CBT combinations shown in **Fig. 2** for each of the four (A, C, T, and G) nucleotides (3×3×4=36 total points) (**Supplementary Table 3**). The different shaped markers correspond to CBT1 (■), CBT2 (●) and CBT3 (▲) barcodes, respectively. Polymerase variants: RPol1 (blue), RPol2 (black) and RPol3 (red). Barcoded DNA templates: CBT1, CBT2 and CBT3.

### Principal component analysis

Our finding that each polymerase variant had a unique set of kinetic parameters opened up the possibility of directly distinguishing them among a variety of polymerase mutants using nanopore sequencing. To evaluate this possibility, we defined three additional kinetic parameters to be used in the principal component analysis (PCA): the tag release rate (TRR) as the frequency of the tag release (**Fig. 3a**, orange double arrow), which is the total time from the last tag capture to the first open channel signal after a nucleotide has been incorporated; tag capture rate (TCR) as the frequency of a tag capture (**Fig. 3a**, blue circle) when a nucleotide is being incorporated into the DNA template; and tag capture dwell (TCD) time as the duration for which the pore conductance is found to be reduced by the presence of a tag. Then, we used PCA on high-quality reads obtained from the sequencing runs for each of the three polymerase variants based on five unique kinetic parameters for each of the four tagged nucleotides (**Supplementary Table 4**). For each polymerase, the PCA-based 2D projections of the kinetic signatures for each polymerase onto the first three principal components showed distinct separation (**Fig. 4** and **Supplementary Movie 1**). Therefore, we demonstrated that polymerase variants could be uniquely identified by using information from multiple kinetic parameters.

**Figure 4.**
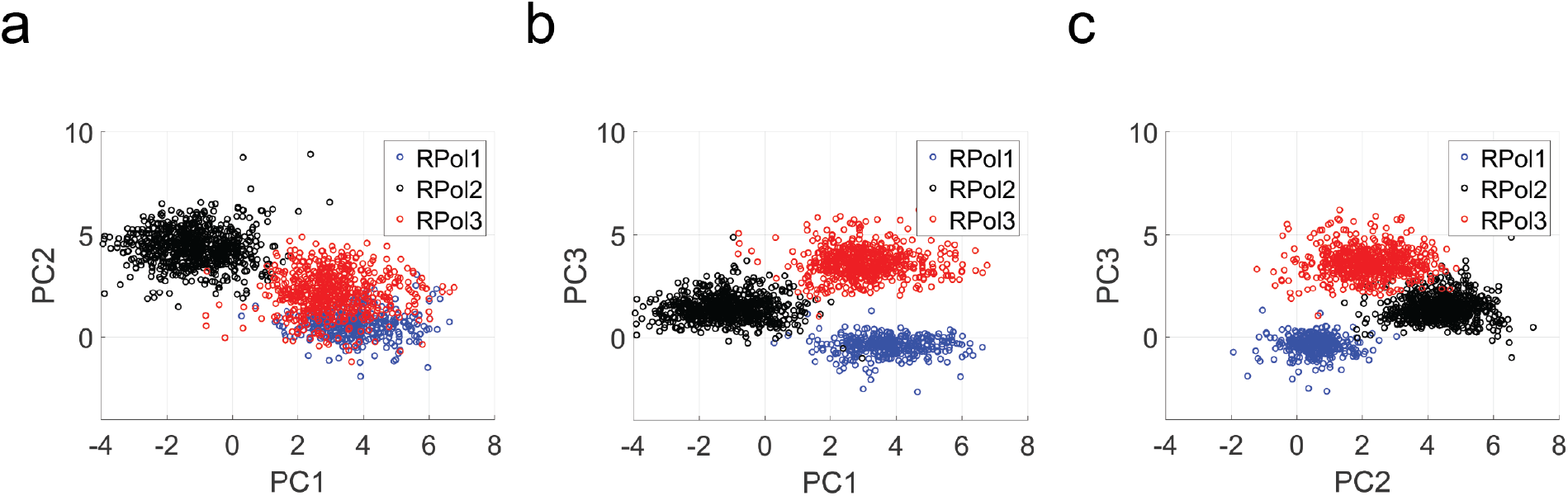
Principal component analysis (PCA) of polymerase variants. Each principal component is a linear combination of 20 parameters (five kinetic properties for each of the four bases) derived from single-molecule tagged nucleotide capture data. The coefficients of these parameters are shown in **Supplementary Table 4**. For each of the three polymerase variants, the PCA-based 2D projections onto the first three (**a**, **b**, and **c**) principal components showed great separation for each of the three polymerase variants. Coordinates on these plots represent *z*-scores computed for the indicated individual principal components. Polymerase variants: RPol1 (blue), RPol2 (black) and RPol3 (red).

### Multiplex polymerase measurement

Our previous experiments established the principle of barcoded-polymerase screening. In practice, one might want to use this approach in a directed evolution scheme to find a polymerase variant with desired kinetic properties. As a proof of principle, we loaded each of the three nanopore-coupled polymerase variants with a unique ssDNA template using a predefined assignment (RPol1:CBT1, RPol2:CBT2 and RPol3:CBT3) in separate template binding reactions. Next, they were pooled in equimolar ratios and inserted into the CMOS chip for sequencing runs. A computationally generated random 51-nt sequence, and a second template, composed of a random 32-mer barcode region with the universal 19-nt flanking priming site, were used as control templates (**Supplementary Fig. 6a**). Utilizing our barcode classification algorithm, on average, we found higher BMPI scores above the threshold value of 0.80 when raw reads were compared to the (correct) templates loaded on the polymerases (**Supplementary Fig. 6b**, Experiments 1 and 3) *versus* two random templates (**Supplementary Fig. 6b**, Experiments 4 and 5). Although, the mean BMPI values were ~0.70 for each RPol:CBT in this pooled experiment, high-confidence barcode identification was still possible as ~67% of the total raw reads (*n* = 418) were identified as any of the three barcodes (**Supplementary Table 5**), which were originally loaded onto the polymerase variants in the pooled 3-plex sequencing experiment.

To explore the potential of multiplexing, we designed 96 synthetic unique barcoded ssDNA templates with the same circular topologies as described for the singleplex experiments (**Supplementary Fig. 1b**). The 32-nt barcoded regions were computationally constructed to serve as unique identifiers by ensuring that the sequence identity calculated by local alignment of any two distinct barcodes was <85% (**Supplementary Fig. 7**). To further test these template designs for high-accuracy barcode identification, we implemented an *in silico* algorithm which sampled 1,000 random high-quality reads from the experiments shown in **Fig. 2**, which were subsequently classified by either comparing them to the experiment-specific (correct) template or to a randomly chosen template from our list of 96 sequences (incorrect template). When the randomly selected high-quality reads were compared to the correct template, the mean BMPI value was 0.85 (**Supplementary Fig. 8**, left). In contrast, when the same reads were compared to randomly selected templates from our list, the average BMPI value shifted below ~0.55 (**Supplementary Fig. 8**, right). This *in silico* test demonstrated the feasibility of a uniquely identifiable polymerase-barcode assignment scheme.

Next, to evaluate these barcoded templates experimentally, we loaded nanopore-coupled RPol2 with these 96 unique CBTs, which were subsequently inserted into a lipid bilayer for sequencing experiments. Then, we used our classifier to analyze the RPol2:CBT1-96 sequencing data for estimating the accuracy with which one could identify each of the loaded CBTs in a single experiment. Each set of high-quality reads obtained was compared to all of the 96 CBTs and a BMPI score was recorded (**Supplementary Fig. 9**). The maximum scoring BMPI value, which was above the 0.80 threshold, identified the most likely barcode candidate for each comparison. Reads with maximum BMPI value less than 0.80 were discarded from downstream analysis. All such classified barcodes were counted and displayed on a histogram (**Supplementary Fig. 10**). Using this classification scheme, we uniquely identified a total of 94 barcodes out of 96 possible (98%) by evaluating 1,067 high-quality raw reads. On average, the individual barcodes were observed at least 20 times during measurements. These observations were randomly distributed as expected by the stochastic nature of pore-polymerase-template assembly and the complex insertion into the lipid bilayer before measurement^28^. Thus, we demonstrated that polymerase-bound barcoded DNA templates could be identified in a 96-plex fashion.

After confirming our capability for the multiplex barcode identification, we further evaluated our method to show multiplex kinetic profiling of multiple polymerases in the same experiment. To test this, we loaded each of the three nanopore-coupled polymerase variants with the first set of 32 templates (RPol1:CBT1-32), the second set from 33 through 64 (RPol2: CBT33-64) and the third set from 65 through 96 templates (RPol3:CBT65-96) from our library of 96 unique CBTs (**Methods**), in separate template binding reactions. Subsequently, they were then mixed in equimolar ratios and inserted into the CMOS chip for sequencing reactions. We used the same barcode classification strategy as for the 96-plex experiments and obtained a randomly distributed frequency histogram in which none of the templates were over/underrepresented for a particular polymerase (**Fig. 5a**). By evaluating 1,958 high-quality raw reads, all of the 96 possible barcodes were identified based on the BMPI cutoff. On average, the individual barcodes were sampled at least 20 times and the observation frequency ranged from 2-68 during measurement. The uneven distribution of the barcode counts (CBT1-32: low, CBT33-64: high, CBT65-96: high) reflects the previously observed processivity differences of the three different polymerase variants (**Fig. 3b**). We also performed three separate control experiments for each of the three prepared complexes to assess the barcode identification specificity in a pooled sequencing reaction. We uniquely identified 20 barcodes (63%) for RPol1:CBT1-32 (number of high-quality raw reads, *n* = 67), and 29 barcodes (90%) for both RPol2:CBT33-64 (*n* = 249) and RPol3:CBT65-96 (*n* = 383) out of the 32 possible barcodes for each set using the same classification scheme as for the single-polymerase, 96-plex experiment (**Fig. 5b**). For RPol1, the individual barcodes were observed at an average frequency of 5, which reflects its slow processivity. Meanwhile, for RPol2 and RPol3 the barcodes were counted at least 10 times on average ranging from 1-28 distinct observations. We showed that barcodes, in their respective set, can be uniquely identified with an average false positive rate of ~13% (**Supplementary Table 6**). Here, we demonstrated that three polymerase variants loaded with multiple different barcoded templates can be identified in a 96-plex fashion.

**Figure 5.**
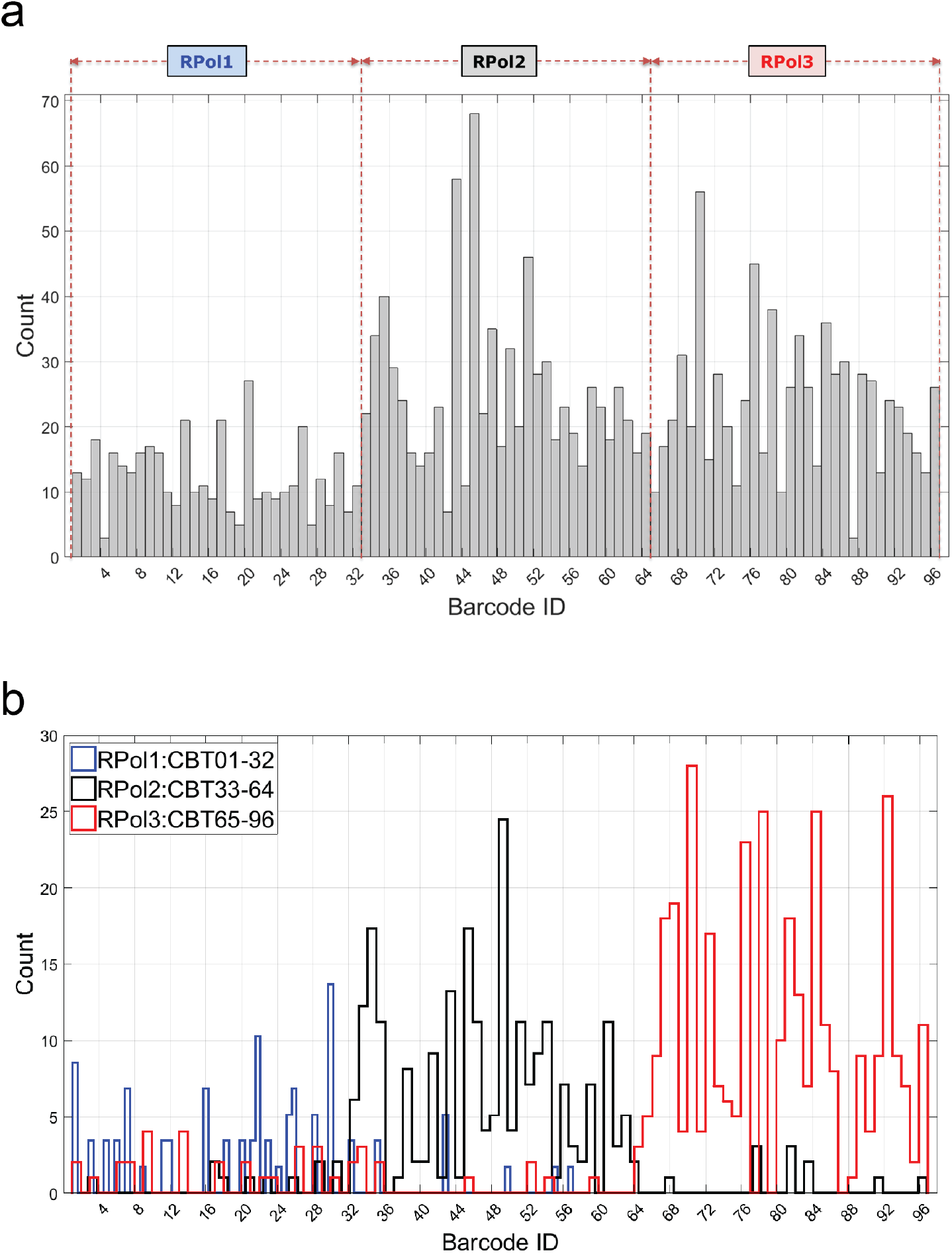
Distribution of barcodes in a multiplex experiment. Circular barcoded templates (CBT) 1-32 were complexed with polymerase variant 1 (RPol1), CBT33-64 with RPol2, and CBT65-96 with RPol3. (**a**) All of the 96 possible barcodes were uniquely identified by the alignment-based classification algorithm (**Methods**). (**b**) Distribution of identified barcodes in individual sequencing experiments for RPol1:CBT1-32 (blue), RPol2:CBT33-64 (black), and RPol3:CBT65-96 (red). Counts are scaled by width of bin for clarity. The expected barcodes are uniquely identified with an average false positive rate of ~13% (**Supplementary Table 6**). Note also the uneven distribution of barcode counts in **a** reflecting the different polymerase processivity as observed in **b**.

Having established the ability of our method to perform multiplexed polymerase identification, we sought to determine how well the barcode sequencing data mapped back to the already determined kinetic properties of a polymerase variant (**Fig. 4**). First, we used PCA on the multiplex sequencing data shown in **Fig. 5a** based on five derived kinetic properties (**Supplementary Table 4**) as before, in which all identified barcodes in each of the barcode sets (CBT1-32, CBT33-64, and CBT65-96, respectively) were accumulated in one group. For each of these barcode groups, the 2D projections of the kinetic properties for each of these barcode groups onto the first two principal components mapped back well (**Supplementary Fig. 11a**), when overlaid with the original PCA clusters derived from the individual singleplex RPol-CBT experiments (**Fig. 4**). Here, the cluster overlay is the measure of the classifier accuracy, which describes how well it can distinguish polymerase variant kinetics based on the barcode sequencing information only. Sequencing data corresponding to the second barcode set (CBT33-64) could not be mapped back well, which could be due to the high false positive rate of barcode identification in that set (**Supplementary Table 6**). On the other hand, sequencing data corresponding to individual barcodes mapped back with high accuracy (**Supplementary Fig. 11b**), which highlights the potential of identifying a single polymerase variant in a multiplex experiment.

Finally, to demonstrate the practical utility of our approach, we sought to scale up our method and apply it towards the kinetic characterization of a library of polymerases to identify variants with different properties. To demonstrate this, we generated a library of 96 polymerase variants (henceforth referred to as LPol) using site-saturation mutagenesis (**Methods**). Stoichiometry of the pore-polymerase conjugates was analyzed by protein gels stained for total protein, which confirmed successful assembly (**Methods** and **Supplementary Fig. 12a**). Next, each of the pore-polymerase conjugates was loaded with a unique template from our library of 96 CBTs (LPol-CBT), thus forming a unique assignment between genotype and phenotype. Subsequently, they were then mixed in equimolar ratios and inserted into the CMOS chip for sequencing reactions. We used the same barcode classification strategy as for the 96-plex experiments and identified top hits based on the number of barcode observations (>3) in the pooled experiment (**Fig. 6a**). By evaluating 1,473 raw reads, 20 polymerase variants were identified to have detectable activity (of the 96 total that were screened) based on the BMPI cutoff. The sparse distribution of the barcode counts might indicate that most polymerase variants in our library had poor processivity, which is below our detection threshold. We identified four polymerase mutants as our top hits: LPol-10 (number of observations, n = 3), LPol-25 (n = 3), LPol-46 (n = 4) and LPol-62 (n = 3) for which the associated barcodes were observed at least 3 times during the experiment. Polymerase function of these top hits was confirmed in an off-chip bulk assay by rolling circle amplification (RCA), which validated their ability to maintain activity after expression and purification (**Methods** and **Supplementary Fig. 12b**). The RCA results also demonstrated that the identified polymerase variants can be used for unidirectional nucleic acid replication to rapidly synthesize multiple copies of circular molecules of DNA. To further characterize the top hits, we calculated the kinetic parameters for each of the four polymerase variants based on the collected sequencing data shown in **Fig. 6a**. We found that the mean FCR was ~2.2 s^−1^, while mean dwell time of the tagged nucleotide captures was computed to be ~0.4 s for LPol-46 (**Fig. 6b** and **Supplementary Table 7**), which indicates that this particular polymerase variant has increased processivity compared to the other three top hits. Similarly, this variant was shown to have one of the highest TRR (~1.5 s^−1^) and TCR (~1.4 s^−1^) among the top hits (**Fig. 6c** and **Supplementary Table 7**), which provides information about the time it takes for the polymerase to be ready to accept the next nucleotide after base incorporation and about the affinity of nucleotide binding to the polymerase during catalysis, respectively. The calculated kinetic parameters, coupled with the results of the off-chip RCA validation experiment (**Supplementary Fig. 12b**), indicate that LPol-46 is a potential candidate to be further evaluated (and evolved) for DNA amplification methods, and biotechnology applications requiring modified nucleotides as active substrates for DNA polymerase, especially for natural or unnatural nucleotides modified on their 5’-phosphate^29,30^. In conclusion, we showed that polymerase variants with a different set of kinetic properties can be uniquely identified by applying our nanopore-based barcode sequencing technique for multiplex library screening. This points towards a future utility of the platform for identifying polymerase variants resulting from a directed evolution scheme with desired kinetic properties, which can be iteratively refined with multiple design (key residue changes to affect kinetic properties), build (site-directed mutagenesis) and test (barcode sequencing of polymerase mutant pool) cycles.

**Figure 6.**
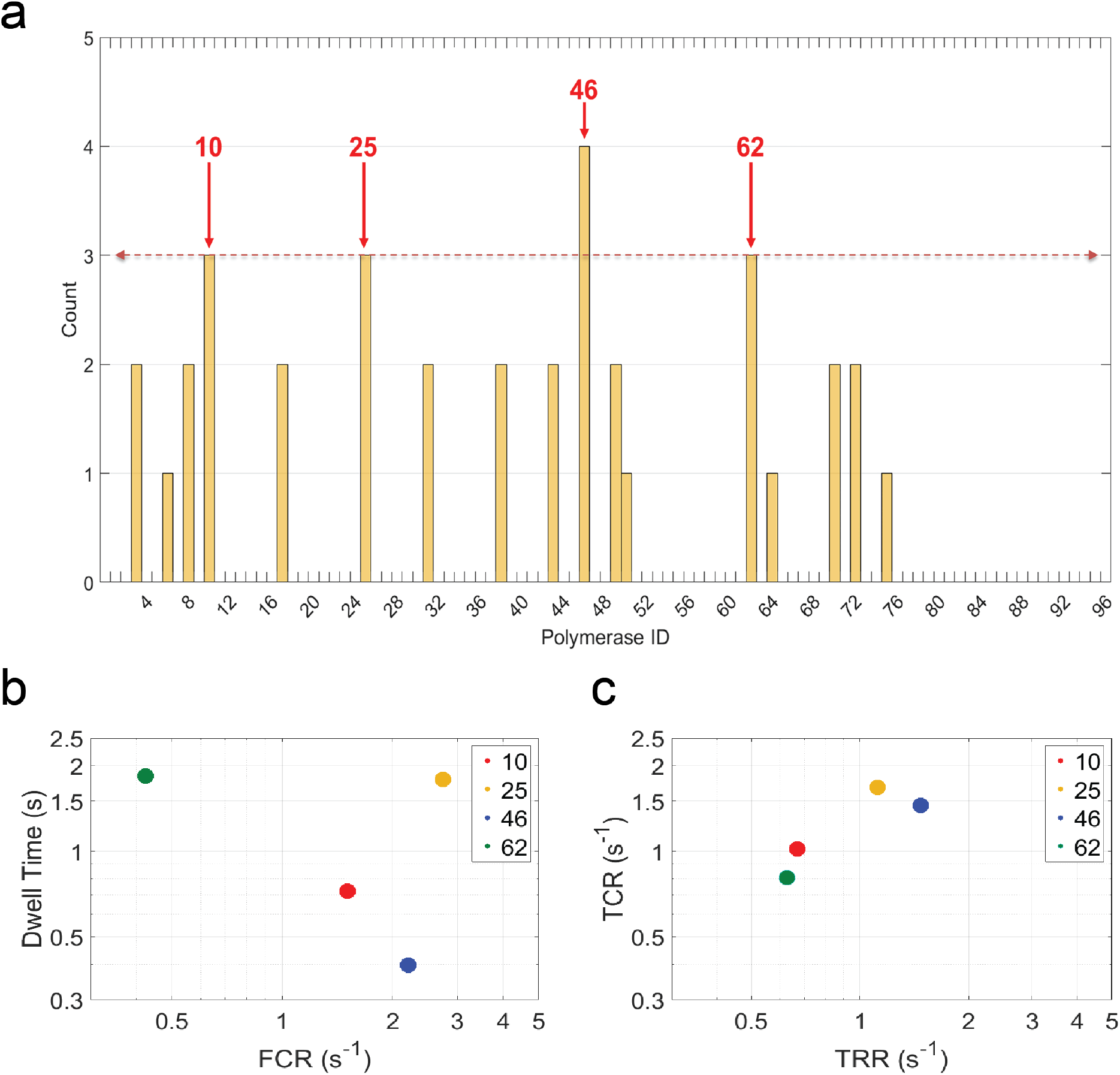
Kinetic screen of a polymerase mutant library. Circular barcoded templates (CBT) were complexed with polymerase variants (LPol) in the library creating a 96 unique LPol-CBT assignment. (**a**) Top hits of polymerase mutants (red arrows: LPol-10, LPol-25, LPol-46 and LPol-62) were identified by the alignment-based classification algorithm based on number of observations (**Methods**). (**b**) Kinetics of top polymerase variant hits. Each marker represents the mean full catalytic rate (FCR) and mean dwell time (t_dwell_) value pair corresponding to each of the LPol-CBT combinations shown in **a** for all of the four (A, C, T, and G) nucleotides. (**c**) Same as **b**, but XY parameters are now for mean tag release rate (TRR) and mean tag capture rate (TCR). The different colored markers correspond to LPol-10 (red), LPol-25 (yellow), LPol-46 (blue) and LPol-62 (green) polymerase variants, respectively.

## Discussion

In our Nanopore-SBS method, polymerase kinetics were monitored in real-time during DNA sequencing. Our results confirmed that we could load polymerases with circular templates and sequence these templates. By enabling repeated interrogation of the same barcoded template, we demonstrated high-sensitivity barcode identification using an alignment-based classification algorithm. These DNA templates also enabled us to distinguish kinetic parameters of different polymerase mutants that have been loaded with unique barcoded templates. We showed high multiplexing potential by performing sequencing reactions on thousands of individually addressable pores on a CMOS chip. The unique kinetic signatures of each polymerase variant, obtained from the barcode sequencing information, permits its discrimination in a pooled experiment. We validated our platform by screening ~100 polymerase variants to identify mutants with different properties, thus showcasing the potential ability of our technology to rapidly screen polymerases at the single-molecule level for applications of biotechnological interest. To our knowledge, this represents the first multiplex study of enzyme kinetics on a nanopore array.

Pacific Biosciences’ approach utilizes fluorescence pulses to identify the nucleotide incorporations, so pulse width and interpulse duration could be used to monitor polymerase kinetics^23^. Alternatively, our screening method could also be implemented on this optical platform after DNA barcoding of a polymerase library and subsequent barcode classification based on single-molecule fluorescent signal derived kinetic parameters similar to the ones derived from the nanopore signal. For this reason, we believe that our method will have broad applicability on different single-molecule platforms.

Future work will focus on developing this nanopore-based platform into a large-scale, multiplex screening tool for DNA polymerases that have user-defined kinetics. As our CMOS-based chip can potentially scale to billions of sensors^31^, this technology could be further extended to a broad spectrum of high-throughput applications in single-molecule enzyme activity or protein-protein interaction studies by correlating the desired molecular event to the observed voltage signature changes through the pore.

## Methods

### DNA template preparation

In the 3-plex experiments, the 51-nt single-stranded DNA (ssDNA) oligonucleotides were computationally designed with a random 32-nt barcode region flanked by a universal 19-nt primer region to uniquely identify each polymerase (**Supplementary Figs. 1d-f**). The synthetic template DNA (IDT) was circularized using CircLigase II (Epicentre), treated with Exonuclease I (NEB) to remove any linear template that was not covalently closed and subsequently column-purified (**Supplementary Fig. 1c**, left panel). As an alternate strategy for circularization, the same sequencing primer was used as a splint to join the ends of the template. Since the primer spanned about ten bases on each end of the template, T4 ligase was then used for ligation and circularization by an overnight incubation at 16 °C (**Supplementary Fig. 1c**, right panel). Unligated linear ssDNA template, excess primer and double-stranded DNA (formed hairpins) were digested with Exonuclease I and III treatment. The resulting primer-annealed circular DNA template was concentrated, desalted and recovered by isopropanol precipitation or by column purification (Zymo Research). The pellet was re-suspended in water and column purified to remove any residual ATP from the previous ligation step. This method yielded high concentrations (>10-fold as compared to the CircLigase method) of the starting template/primer complex, and hence the template:polymerase:pore ratio in the final reaction could be scaled up accordingly. Therefore, this method was used for subsequent circularization experiments. The primer (5’-ATTTTAGCCAGAGTGGGGA-3’) was then annealed to the circularized barcoded template by heating to 95 °C for 3 min followed by cooling to 20 °C at a rate of 0.1 °C/s.

For the multiplex experiments, a set of 96 unique barcoded ssDNA templates were computationally designed and ordered (IDT). The 32-nt barcoded regions were constructed such that when any one of the templates was locally aligned to all other templates in the full set, the calculated sequence identity was always <85% to generate unique identifiers (**Supplementary Fig. 7**). They were then either divided into three individual sets (set 1 = CBT 1 through 32; set 2 = CBT 33 through 65 and set 3 = CBT 66 through 96), wherein each set consisted of 32 templates, or all 96 templates were pooled together. Finally, each of these sets of 32 or the 96 pooled templates were circularized and primer-annealed for subsequent reactions.

### Preparation of polymerase and tagged nucleotides

Clostridium phage ϕCPV4 DNA polymerase (GenBank ID AFH27113.1) was used as wild-type. For the RPols, specific mutations were introduced to the DNA polymerase gene by site-directed mutagenesis (Roche Sequencing Solutions) to enhance the kinetic properties of the polymerase utilizing polynucleotide tagged nucleotides to approach native nucleotide incorporation characteristics^32^. For the polymerase screen (LPols), a randomized site-saturation mutagenesis library was designed based on homology alignment with ϕ29 DNA polymerase (PDB ID 2PYL). Five target residues (K237, E323, K325, K453, K455) were randomly chosen in the N-terminal and palm domains, to minimally impact nucleotide binding, to generate all 19 different single amino acid substitutions resulting in 96 different mutants. Each mutant was individually transformed to BL21(DE3) cells for downstream protein expression and purification steps (Thermo Fisher). Tagged nucleotides were synthesized as described previously^28^. Briefly, click chemistry was utilized to link the 5’-phosphate and the polynucleotide tag interspersed with a variety of chemical moieties for each of the four nucleotides [dA6P-Cy3-dT_30_-C3, dC6P-Cy3-dT_5_-(BHEB)-dT_24_-C3, dT6P-Cy3-dT_4_-(N3CET)_3_-dT_23_-C3, dG6P-dT_6_-(dTmp)_6_-dT_19_-C3, where abbreviations are defined as “BHEB”=bis-hydroxyethylbenzene; “N3CET”=3-N-cyanoethyl-dT amidite; “dTmp” = thymidine methyl phosphonate].

### Protein expression and purification

LB medium supplemented with 12 μg/mL kanamycin was inoculated from a glycerol stock of BL21(DE3) cells and incubated overnight at 37 °C and 800 rpm in a 96-well plate format. 30 μL of the overnight cultures were transferred into 1.25 mL of fresh LB media in new 96-well plates. Growth of cultures was continuously monitored by taking OD_600_ measurements every **~**30 min and at OD ~0.6 protein expression was induced by supplementing the cultures with 1 mM isopropyl β-D-1-thiogalactopyranoside and incubating overnight at 15 °C and 800 rpm. Cells were harvested by centrifugation for 20 minutes at 4 °C and 3500 g. The protein library was purified using a His-Tagged 96-well plate cartridge (Clontech) and buffer exchanged into 50 mM Tris-HCl, pH 7.5, 150 mM NaCl, 5 mM tris (2-carboxyethyl) phosphine (TCEP), 8% (w/v) trehalose, and 0.01% (v/v) Tween 20 using spin desalting plates (ThermoFisher) per manufacturer’s instructions. Concentration of the purified protein was determined with a reducing agent compatible micro BCA kit (Thermo Fisher) as per manufacturer’s protocol and averaged to ~200μg/mL. Select polymerase variants were characterized by 4-12% Bis-Tris protein gels (Thermo Fisher) to confirm the presence of polymerase and pore conjugates (**Supplementary Fig. 12a**).

### Confirmation of polymerase function

Polymerase function was determined with a real-time rolling circle amplification (RCA) assay (**Supplementary Fig. 12b**). In brief, the set of 96 CBTs, which was utilized in the multiplex experiments, was used as pooled templates. Wild type ϕ29 was purchased from NEB as positive control. RCA was performed at 30 °C in a Mastercycler Realplex qPCR (Eppendorf) for 60 min in 20 μL reactions. Each reaction contained 1x ϕ29 reaction buffer (NEB), 100 nM pooled CBTs, 10 nM primer (5’-ATTTTAGCCAGAGTGGGGA-3’), 0.3 mM dNTP, 1x SYBR ssDNA dye (Thermo Fisher), and 1 μL of protein sample.

### Pore-polymerase-template complex formation

Purified polymerase and the desired template were bound to a αHL pore (*via* SpyCather-SpyTag chemistry) by incubating 0.1 M polymerase and 0.1 M of primer-annealed circularized DNA template per 0.1 M of 1:6 pore overnight at 4 °C to form the pore-polymerase-template assembly as described previously^28^. For the spike-in experiments to test template switching, 2-fold molar excess of the desired template was first incubated with the polymerase, and then with the 1:6 pore overnight, before loading onto the chip.

### Nanopore experiments

Synthetic lipid 1,2-di-*O*-phytanyl-sn-glycero-3-phosphocholine (Avanti) was diluted in tridecane (Sigma-Aldrich) to a final concentration of 15 mg/mL. A planar lipid bilayer was formed on the CMOS chip (Roche Sequencing Solutions) surface containing an array of 32,768 electrodes as described previously^28^. The electrodes were arranged in a rectangular array of 64 rows and 128 columns, with 2.4 μm spacing between rows and 2.6 μm spacing between columns at a pitch of 16μm. Sequencing experiments were performed in asymmetric conditions. The *cis* compartment was filled with a buffer containing 300 mM KGlu, 3 mM MgCl_2_, 10 mM LiCl, 5 mM TCEP and 20 mM HEPES pH 8.0 and the *trans* compartment with 380 mM KGlu, 3 mM MgCl_2_ and 20 mM HEPES pH 8.0, in which MgCl_2_ is a catalytic cation source during the polymerase extension reaction to initiate and sustain sequential nucleotide additions along the template DNA. In our buffers, KGlu was used to increase DNA-protein interactions^33^ and LiCl to slow down the DNA translocation^34^ of the tag through the pore. Purified pore-polymerase-template conjugates were diluted in buffer to a final concentration of 2 nM. After pumping a 10 μL aliquot to the *cis* compartment, single pores were embedded in the planar lipid bilayer that separates the two compartments each containing ~5 μL of buffer solution. Experiments were conducted at 20 °C with 10 μM tagged nucleotides added to the *cis* well. Typically, ~70% of the total pores contained a functional polymerase complex. During the various experimental steps, a precision syringe pump (Tecan) was utilized in an automated fashion to deliver reagents into the microfluidic chamber of the CMOS chip at a flow rate of 1 μL/s. Software control was implemented in Python, which interfaced with the pump *via* an RS 232 communication protocol.

### Electrical recording and data acqusition

To demonstrate the data acquisition technique, we evaluated an open channel transition to a tag capture state (**Supplementary Fig. 14a**, red box). First, a positive voltage bias (V_min_=+220 mV) (**Supplementary Fig. 14b**, middle panel) was applied across the nanopore in response to the reset signal, which forced the tag to be captured in the pore (**Supplementary Fig. 14b**, top panel: state 3). The reset signal was kept high for a short time period (t_1_=200 μs) during which the capacitor was charged. Then, the reset signal was kept low (therefore V_min_=0 mV) for a longer time period (t_2_=8 ms), so that it was discharged, and the rate of decay was recorded with a sampling rate of 2 kHz – resulting in a stream of ADC values taken every ~0.5 ms (**Supplementary Fig. 14b**, lower panel: state 3). This last voltage command sequence was defined as the “read” period. Immediately after, the same procedure was repeated, but now with a negative voltage bias (V_min_=-10 mV for t_1_*=200 μs) (**Supplementary Fig. 14b**, middle panel) which forced the captured tag to be ejected from the pore (**Supplementary Fig. 14b**, top panel: state 4). The subsequent discharge stage (V_min_=0 mV) was defined as the “eject” period with time duration of t_2_*=12 ms, which was sampled by ADC with the same frequency as in the “read” period (**Supplementary Fig. 14b**, lower panel: state 3). The “read” and “eject” commands were continuously alternated during a sequencing experiment.

### Nanopore signal normalization

Normalization can allow compensating for changes in the electrical properties of an individual sequencing element (sequencing unit) in the nanopore array. The voltage across the nanopore corresponds to the voltage difference (*V*) between the electrode pairs (**Fig. 1a**, components **1** and **3**), which is a function of the capacitance of the lipid bilayer (*C*) according to equation of *C=q/V*, given the same amount of charge *q*. The membrane capacitance may vary over time due to the physical changes (deformation) in the bilayer area or thickness and thus changing the voltage gain, which is referred to as voltage drift. Additionally, charge can build up in the sequencing unit, which is attributed to the differences in the charge transfer between “read” periods and “eject” periods (**Supplementary Fig. 14**), which is referred to as baseline shift. These are the two primary causes of errors in real sequencing systems.

To correct for these effects, ADC values measured during sequencing were normalized to provide greater base-calling accuracy. Data points recorded during the “read” periods (**Supplementary Fig. 14b**, states 1 & 3; **Supplementary Fig. 14c**, states 3 & 5) were used in the normalization algorithm while points collected during the “eject” periods (**Supplementary Fig. 14b**, states 2 & 4; **Supplementary Fig. 14c**, states 4 & 6) were omitted as they did not contain information related to tag captures. In general, the reduction of pore conductance during a “read” period indicated that a tag was captured in the pore. Whenever the ADC value deflected below 80% of the open channel level (~150 ADC counts) for a particular sampling point, it was defined as a blocked state. Similarly, unblocked states were sampling points in the “read” periods for which the pore conductance stayed at the open channel level. These states indicated that no tag is captured in the pore during that time. Data were recorded with a sampling rate of 2 kHz; thus 16 distinct ADC values were measured during the “read” period (t_2_ = 8 ms). ADC values acquired during a “read” period were normalized by dividing each measured data point by the averaged ADC values measured in continuous unblocked states. By utilizing this normalization technique, the dynamic range of the raw ADC measurements was rescaled to a normalized range of [0,1]. More specifically, to calculate the FOCS *s* of a sampling point *i* in a “read” period, we took an average of the ADC values *s*_*j*_ measured in continuous unblocked states during the same “read” period (*j* ∈ 1: *N*, where *N* is the number of unblocked states), and divided the measured ADC value of each sampling point by this local mean ADC value:

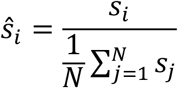

The median was determined for the normalized ADC values in the “read” period and was reported as FOCS (at mid-period point), identifying the different states of the nanopore. Tagged nucleotide incorporations (TNI), which were indicated by continuous “read” periods in the blocked state, were base-called by a similar classification algorithm described in our previous work^28^. Briefly, TNIs were differentiated from background events by requiring their continuous “read” cycles in the blocked state to be greater than 2. Then, we assigned base calls (see **Fig. 1b**) to all such TNIs by comparing their mean FOCS to the four FOCS bands bounded by the first and third quartile mean values (lower/upper bounds) of the FOCS corresponding to a particular TNI (A,C,T,G) respectively (**Supplementary Figs. 2b, 15**).

### Principal component analysis

Standard principal component analysis was carried out using the *pca* function from the Statistics and Machine Learning Toolbox of MATLAB (2017a, MathWorks, Natick, MA). Input variables were scaled to have zero mean and unit variance and the resulting first, second and third principal component were determined from the entire dataset (**Supplementary Table 4**). To generate the principal component scatter plot (**Fig. 4**), all of the sequencing data for each polymerase variant were first projected onto these first three principal components. These values were then converted into a *z* score by centering and scaling of all data points for each principal component. For each polymerase variant, **Supplementary Movie 1** shows the 3D separation between projections of the kinetic parameters for each polymerase variant onto the first three principal components.

### Classification of barcodes

Voltage signal events were converted to raw reads using a commercial probabilistic base-calling algorithm (version 2.9.2, Roche Sequencing Solutions, Santa Clara, CA). Raw reads, with read lengths greater than one full barcode iteration (51 bp), were then fed as input to a Smith-Waterman (SW) alignment-based barcode classification algorithm, which assigns a BMPI value to that read. More specifically, the first step was to classify the different regions in the raw circular reads into barcode reads. This was achieved by locally aligning the raw read sequence to the known concatenated barcode sequence, where the concatenation multiplier (CM) is calculated by the following formula:

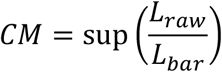

where *L*_*raw*_ is the length of raw read; *L*_*bar*_ is the length of barcode and *CM* is an integer. Once all barcode iteration boundaries were identified, we utilized the *multialign* function from the Bioinformatics Toolbox of MATLAB (2017a, MathWorks, Natick, MA) to perform a progressive multiple alignment of the repeated barcode sequences. Next, we generated the consensus sequence of these multiple aligned reads using *seqconsensus*, which was subsequently locally aligned to all potential barcodes in the experimental set, if the consensus sequence length was at least 10 bp. Finally, the maximum scoring (SW) alignment identified the most likely barcode candidate, which was evaluated based on the particular input sequence. This score was defined as the BMPI and is used to measure the barcode identification probability with possible range of [0,1], where 0 means total mismatch and 1 denotes a total match. For all alignments, homopolymer sequences in the template, and repeated base calls of the same nucleotide in the raw sequencing reads were considered a single base.

### High-quality raw reads

To filter out high-quality raw reads for barcode identification, we have generated the cumulative BMPI of all three polymerase variants as a function of full barcode iterations. We have observed that in general as the read length increases the BMPI of the barcodes asymptotically increases up until 10, 14 and 20 iterations for RPol1, RPol2, and RPol3 respectively. As a conservative approach, we have considered raw reads with at most 10 full iterations for barcode identification, while the rest of the other sequences where discarded in the downstream analysis pipeline (**Supplementary Fig. 3**).

## Acknowledgements

We acknowledge John Aach, Benjamin Boettner, Hani Sallum and Richard Novak for helpful discussions and comments on the manuscript. This work was supported by grants from the National Human Genome Research Institute (R01 HG007415), the National Science Foundation (MCB-1445570), and Roche Sequencing Solutions.

## Author contributions

M.P., S.P, P.B.S., and G.M.C designed the study. M.P., S.P., P.B.S. performed experiments, M.P. analyzed data, and M.P., S.P. wrote the manuscript. F.V., J.N., D.J.W., A.A., T.C., D.G., H.F., S.S., J.P., A.T., C.A., C.S., C.M., P.S., D.D., E.T., E.A., I.L., M.T., S.K. and A.Q. contributed new reagents/analytic tools. M.P. wrote custom MATLAB analysis software. C.W.F., S.R., and G.M.C. discussed results and commented on the manuscript.

## Data availability

The data that support the findings of this study are available from the corresponding author upon reasonable request.

## Code availability

Custom codes for data analysis are available *via* https://github.com/pirimidi/Enzymatic-Profiling.

