## Supplementary Information for "Multiplex single-molecule kinetics of nanopore-coupled polymerases"

a

**CBT1**

5′ – GGC TAA AAT CCT TAA TTA ACC CTT TTC CCT TTT GAA TTC CCT CCC CAC TCT – 3'

**CBT2**

5′ – GGC TAA AAT TGG TAC ACG GGT CCT TAC GGT TTC TTA CAC TCT CCC CAC TCT – 3'

**CBT3**

5′ – GGC TAA AAT GGA ATT AAT TGG GAA AAG GGA AAA CTT AAG GGT CCC CAC TCT – 3'

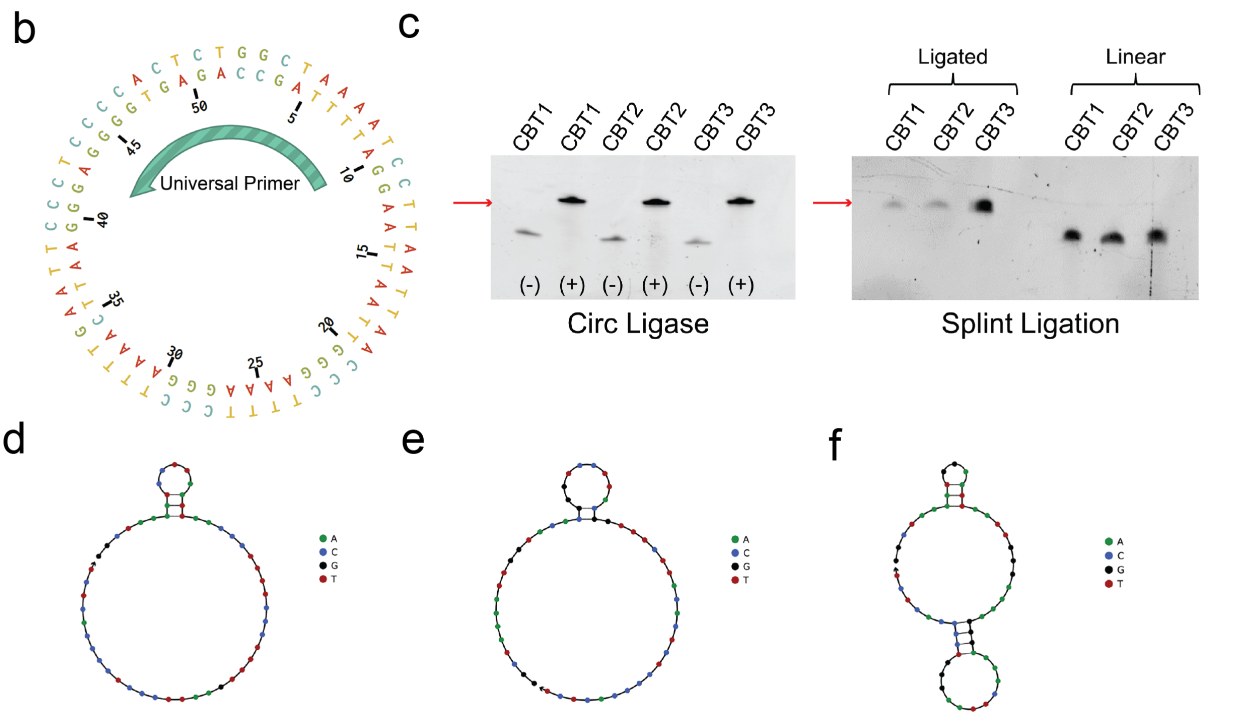


**Supplementary Figure 1** Barcode design. (**a**) Linear sequence of the three 51-bp barcoded ssDNA templates (CBT). The 19-bp universal priming region, used in template circularization and in the sequencing reaction, is highlighted in yellow. (**b**) Representative circular map of a CBT, after a full sequencing iteration (now dsDNA), is shown with a 19-bp priming site (cyan). (**c**) Denaturing urea gel showing the three different linear ssDNA templates that have been circularized using CircLigase II (Epicentre) [left panel] or splint ligation using T4 ligase [right panel]. Closed circular DNA migrates slower, thus appears higher (red arrow) than linear DNA on the gel. (**d-f**) Predicted secondary structures of the three 51-bp ssDNA templates used as barcodes. Minimum-free energy structures are predicted at 37 °C by NUPACK[1,2]. Nucleotides are shown in standard Sanger colors.

a

b

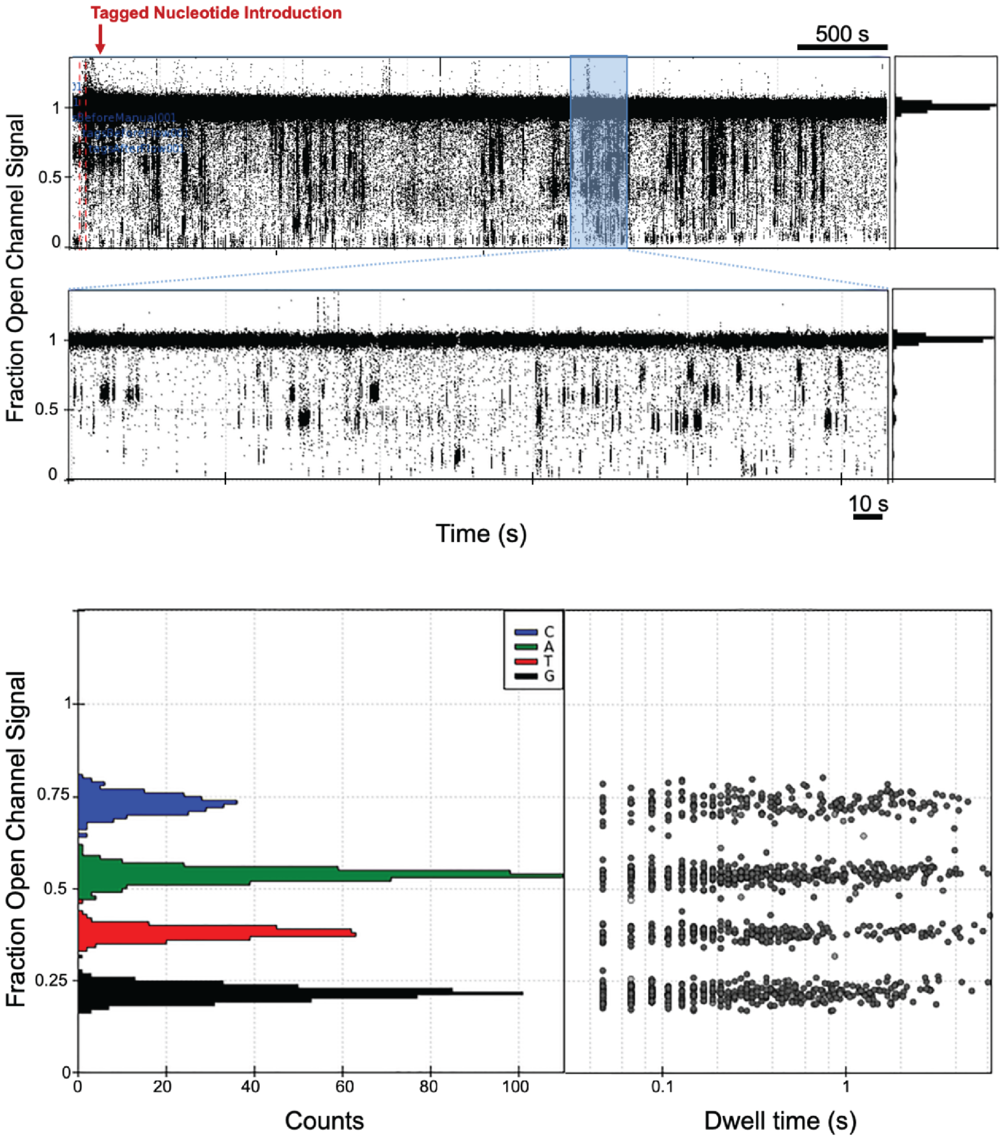


**Supplementary Figure 2** Typical tag capture signal with a single pore-polymerase-template complex RPol1:CBT1 on chip. (**a**) This sequencing experiment demonstrates that the barcoded complex readily inserts into the lipid bilayer and shows template-specific capture signal for over 4,000 seconds [top panel]. The bottom panel shows a zoomed region (300 s) of the same trace with the tag capture signal pronounced. Histogram of fraction open channel signal (FOCS) is shown in the right sub-panels. Dominant peak corresponds to the open channel signal (FOCS=1), while the four minor peaks represent signal associated with each tagged nucleotide capture. (**b**) FOCS histogram of base captures for each tagged nucleotide during a typical barcode sequencing experiment.

**Supplementary Table 1** Number of quality raw reads analyzed for each polymerase-barcode combination.

| | Barcode | | |
|---|---|---|---|
| **Polymerase** | CBT1 | CBT2 | CBT3 |
| RPol1 | 363 | 649 | 407 |
| RPol2 | 461 | 332 | 479 |
| RPol3 | 459 | 490 | 399 |

Polymerase variants: RPol1, RPol2 and RPol3. Barcoded DNA templates: CBT1, CBT2 and CBT3.

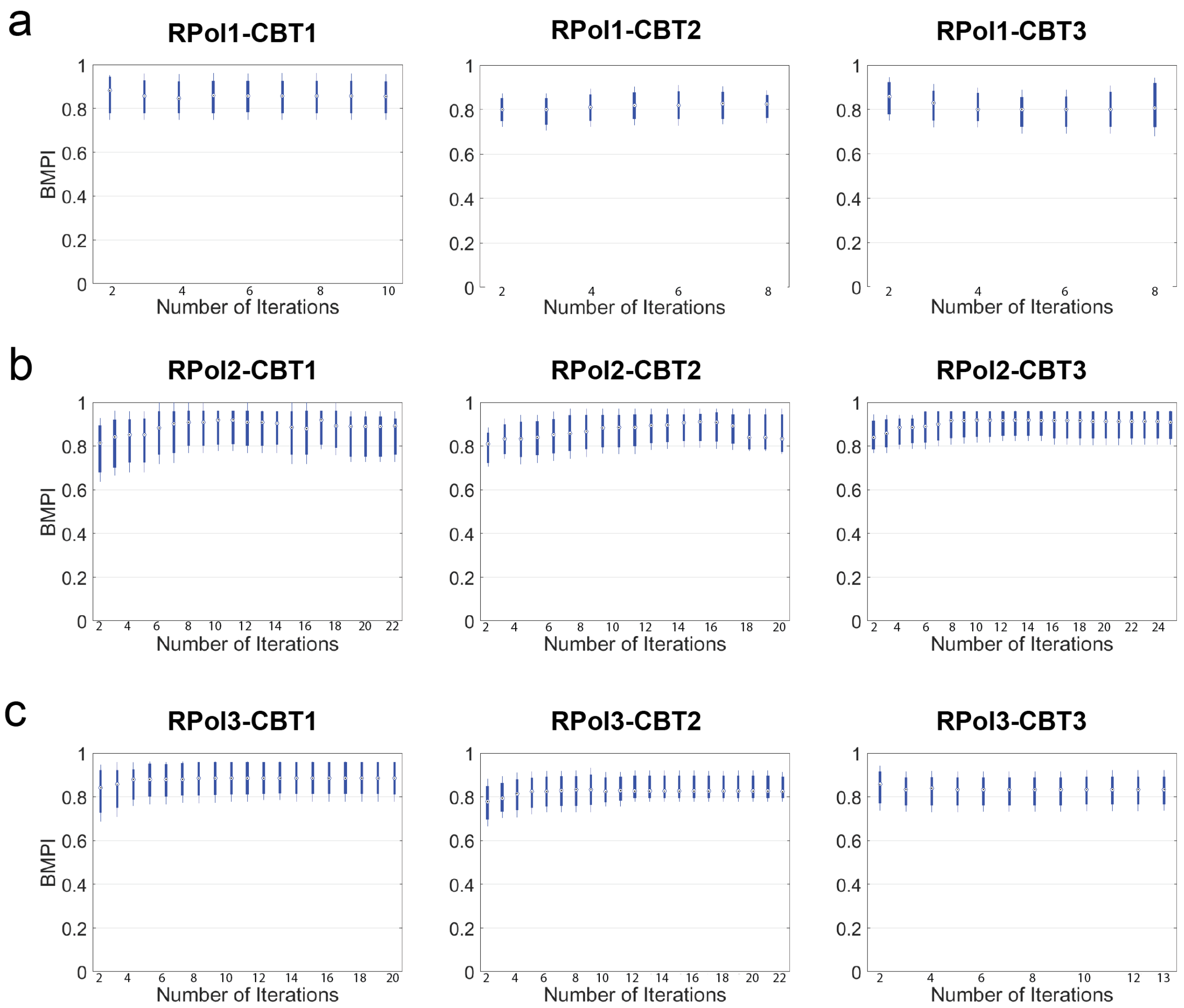


**Supplementary Figure 3** Dependency of barcode identification probability on read length. Quality raw reads are cumulatively grouped together with at least *N* iterations for each polymerase-barcode sequencing experiment. The barcode match probability index (BMPI) values logarithmically increase up to 10 full barcode iterations (~500 bp) then reach an asymptotic saturation point.

**Supplementary Table 2** Percent probability of misidentifying the known barcode as another barcode when the mean BMPI value, computed by the barcode classifier, is greater than 0.80.

| BMPI Values | RPol1 | RPol2 | RPol3 |
|---|---|---|---|
| Total Raw Reads | 1686 | 2278 | 2214 |
| Raw Reads with BMPI > 0.80 | 27 | 15 | 48 |
| % of Misidentified Barcodes | 1.60 | 0.66 | 2.17 |

BMPI values, calculated when the raw reads were compared to the incorrect barcodes, were lumped together into the same dataset for each of the three polymerase variants (RPol1, RPol2, RPol3) loaded with a known template.

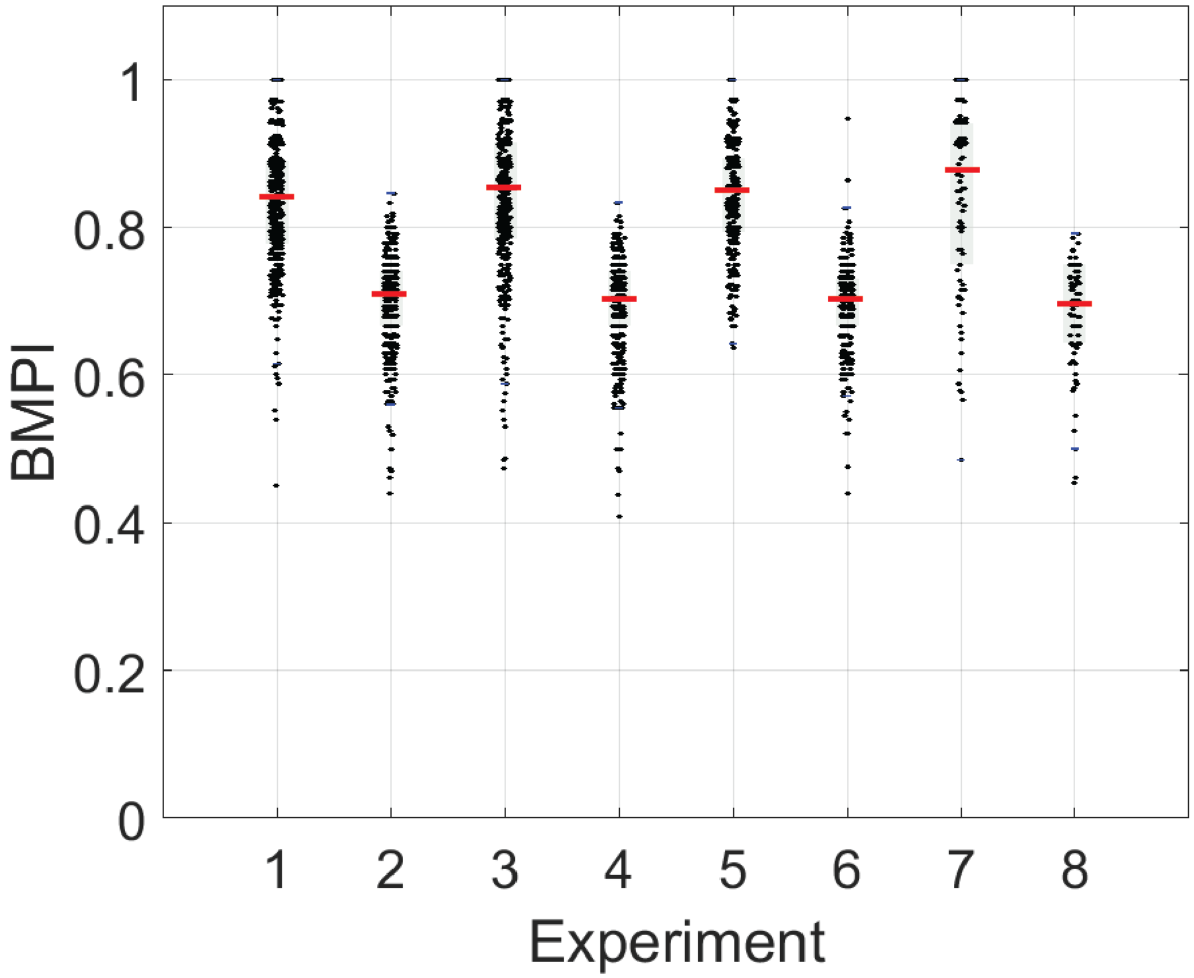


**Supplementary Figure 4** Barcoded DNA template replacement. (**1**) Barcode match probability index (BMPI) values of pore-polymerase-template complex RPol2-CBT2 when comparing to the expected barcode, CBT2. Number of quality raw reads (*N*) = 612. (**2**) When comparing to an incorrect barcode (CBT1), the sequencing accuracy dramatically dropped. (**3**) The presence of a non-complexed barcode (CBT1), immediately spiked in after the nanopore-polymerase-barcode complexing, did not indicate barcode replacement. (**4**) When reads in **3** were compared to the incorrect barcode (CBT1), we observed a similar result as for our control case in **2**. (**5**) Even, after an overnight incubation with a second barcode (CBT1), we observed no barcode switching. (**6**) We also tested on-chip barcode replacement, when we spiked in the second barcode (CBT1) along with the tagged nucleotides after pore insertion. (**7**,**8**) These results indicated that the polymerase variants are uniquely labeled with their respective barcodes and are not replaced in an experiment.

**Supplementary Table 3** Statistical details.

| FCR (μ ± SEM) | | | | | | | | | |
|---|---|---|---|---|---|---|---|---|---|
| | RPol1 | | | RPol2 | | | RPol3 | | |
| | CBT1 | CBT2 | CBT3 | CBT1 | CBT2 | CBT3 | CBT1 | CBT2 | CBT3 |
| A | 0.66 ± 0.01 | 0.57 ± 0.02 | 0.73 ± 0.03 | 1.35 ± 0.03 | 1.30 ± 0.02 | 1.60 ± 0.03 | 1.95 ± 0.03 | 1.89 ± 0.03 | 2.05 ± 0.05 |
| C | 0.55 ± 0.02 | 0.56 ± 0.02 | 0.57 ± 0.02 | 1.48 ± 0.06 | 1.89 ± 0.04 | 1.90 ± 0.04 | 2.27 ± 0.07 | 2.99 ± 0.07 | 2.60 ± 0.08 |
| G | 0.83 ± 0.02 | 0.66 ± 0.01 | 0.86 ± 0.05 | 1.39 ± 0.04 | 1.44 ± 0.03 | 1.46 ± 0.03 | 2.10 ± 0.03 | 2.02 ± 0.04 | 2.02 ± 0.06 |
| T | 0.81 ± 0.02 | 0.66 ± 0.02 | 0.89 ± 0.03 | 1.62 ± 0.05 | 1.42 ± 0.04 | 1.77 ± 0.03 | 2.19 ± 0.04 | 1.80 ± 0.04 | 1.96 ± 0.06 |

| $t_{dwell}$ (μ ± SEM) | | | | | | | | | |
|---|---|---|---|---|---|---|---|---|---|
| | RPol1 | | | RPol2 | | | RPol3 | | |
| | CBT1 | CBT2 | CBT3 | CBT1 | CBT2 | CBT3 | CBT1 | CBT2 | CBT3 |
| A | 1.25 ± 0.02 | 1.15 ± 0.04 | 0.98 ± 0.05 | 0.79 ± 0.01 | 0.76 ± 0.02 | 0.67 ± 0.01 | 0.55 ± 0.01 | 0.50 ± 0.01 | 0.50 ± 0.01 |
| C | 1.57 ± 0.05 | 1.63 ± 0.04 | 1.79 ± 0.06 | 0.82 ± 0.04 | 0.77 ± 0.02 | 0.83 ± 0.02 | 0.52 ± 0.02 | 0.52± 0.01 | 0.61 ± 0.01 |
| G | 1.21 ± 0.02 | 1.19 ± 0.02 | 1.13 ± 0.05 | 0.81 ± 0.02 | 0.66 ± 0.02 | 0.68 ± 0.02 | 0.55 ± 0.01 | 0.46 ± 0.01 | 0.49 ± 0.01 |
| T | 0.94 ± 0.02 | 0.95 ± 0.03 | 0.94 ± 0.04 | 0.71 ± 0.02 | 0.71 ± 0.02 | 0.71 ± 0.01 | 0.54 ± 0.01 | 0.55 ± 0.01 | 0.63 ± 0.01 |
| # | 214 | 115 | 60 | 172 | 271 | 306 | 243 | 224 | 142 |

For each of the RPol:CBT combinations, mean (μ) and standard error of the mean (SEM) values corresponding to the full catalytic rate (FCR) and dwell time ($t_{dwell}$) for each of the four (A, C, G, and T) nucleotides are shown. #: number of quality reads used in the calculations. The unit of FCR and $t_{dwell}$ is $s^{-1}$ and s, respectively.

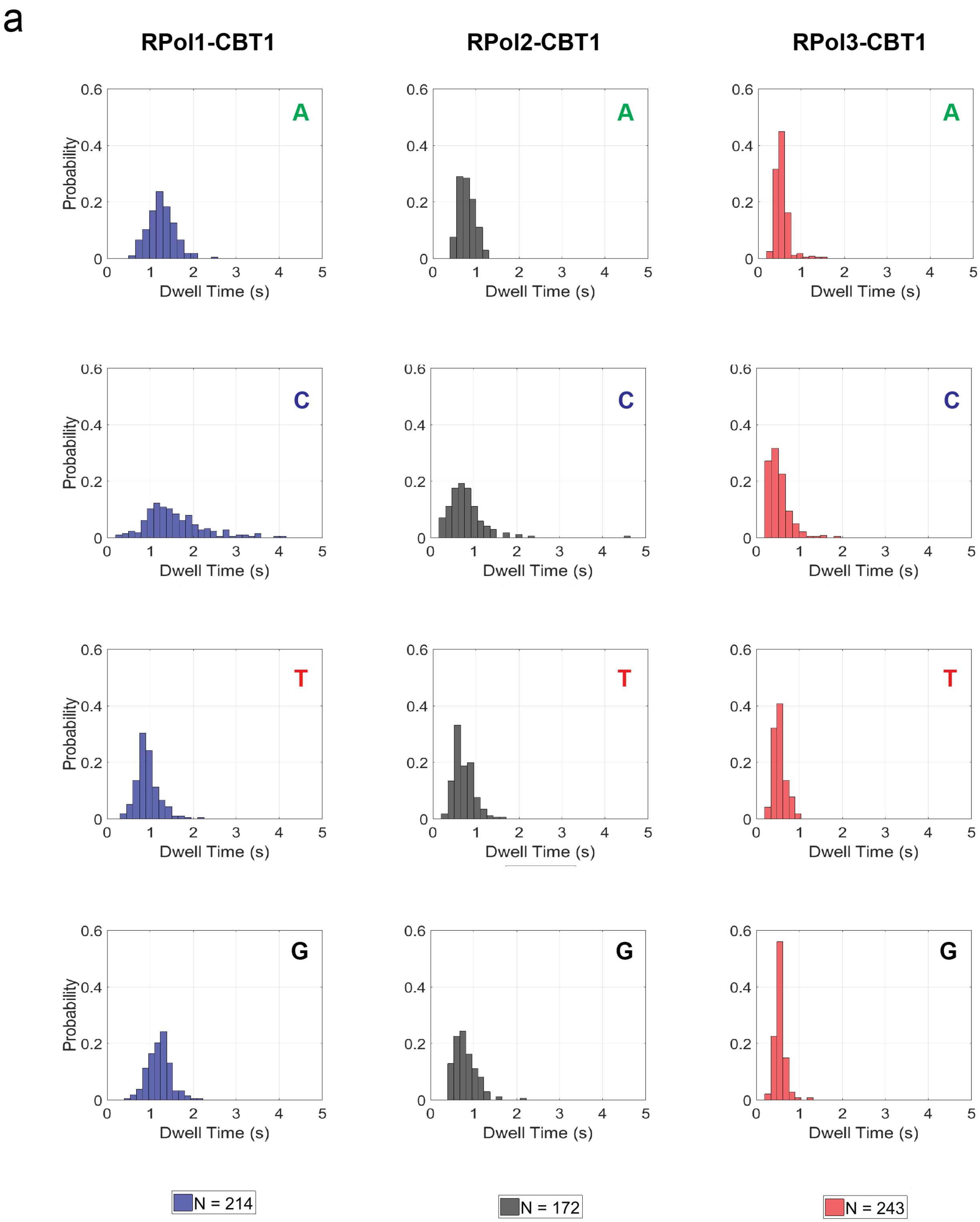
a
RPol1-CBT1
RPol2-CBT1
RPol3-CBT1
A
C
T
G
Probability
Dwell Time (s)
N = 214
N = 172
N = 243

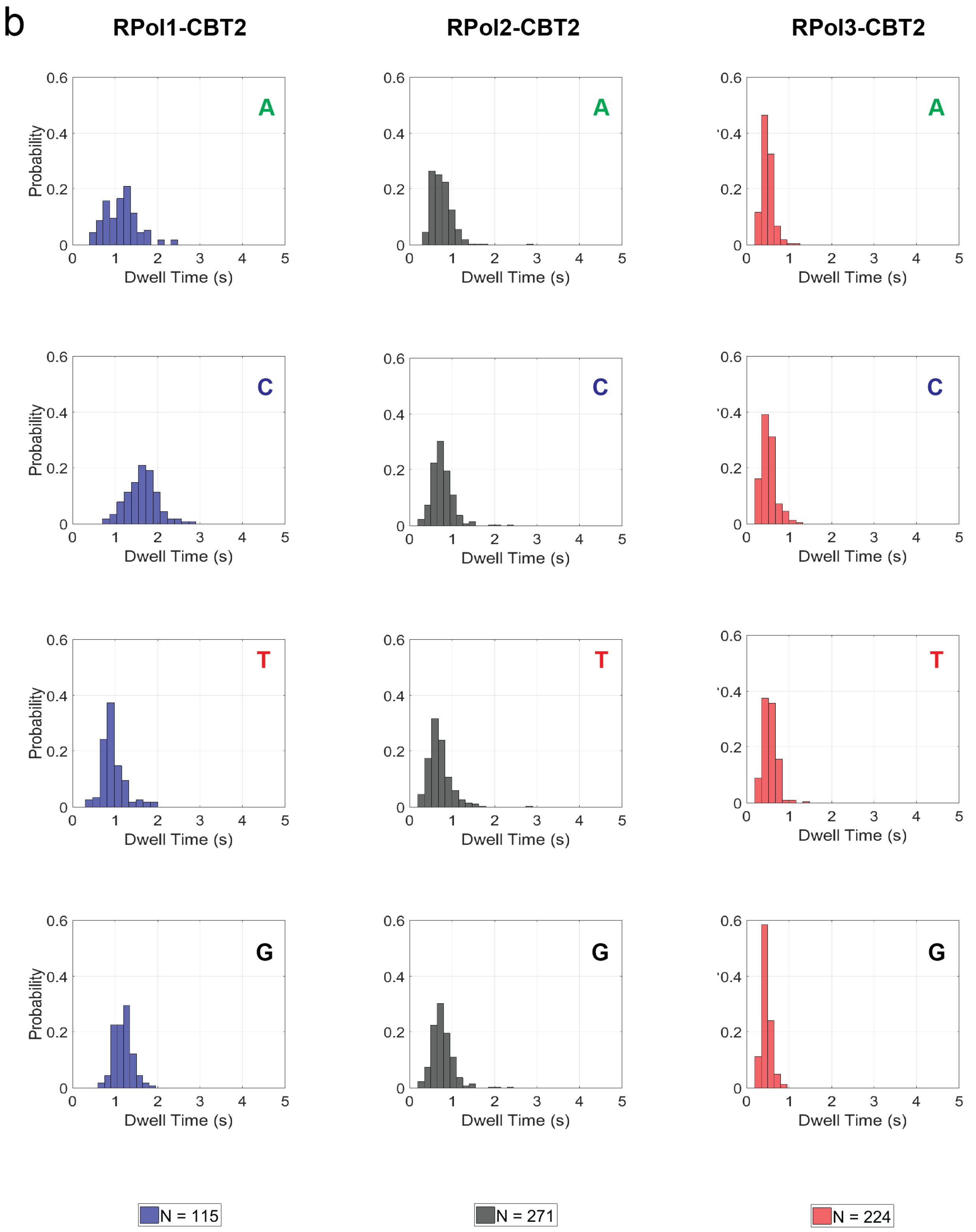
b
RPol1-CBT2
RPol2-CBT2
RPol3-CBT2
A
C
T
G
Probability
Dwell Time (s)
N = 115
N = 271
N = 224

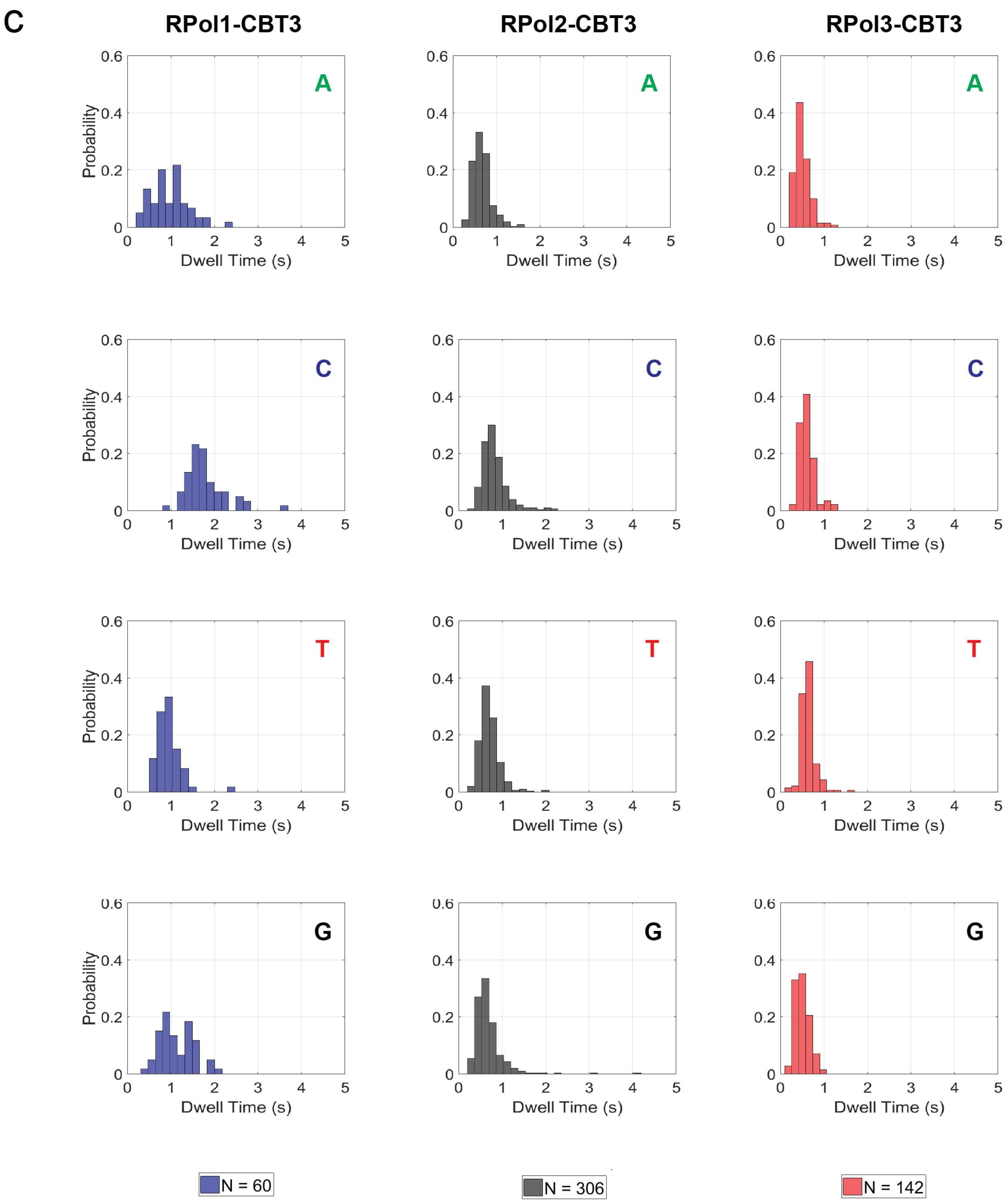
C
RPol1-CBT3
RPol2-CBT3
RPol3-CBT3
A
C
T
G
Probability
Dwell Time (s)
N = 60
N = 306
N = 142

**Supplementary Figure 5** Dwell time histograms. Mean dwell time distribution of each of the four tagged nucleotides for each of the three polymerases (RPol1-3) loaded with (**a**) circular barcoded template (CBT) 1, (**b**) CBT2 and (**c**) CBT3. The distributions do not differ from barcode to barcode, which demonstrates that dwell time, a polymerase-associated kinetic property, is independent of barcode choice. On the other hand, for each polymerase variant, the mean dwell time is different: centers around 1.2 s, 0.7 s and 0.5 s, respectively. Thus, dwell time is a kinetic property that could be used to distinguish polymerase variants.

**Supplementary Table 4a** Principal component analysis coefficients for RPol1.

| Parameters | PC1 | PC2 | PC3 |
|---|---|---|---|
| A-FCR | -0.0206 | -0.9631 | -0.4112 |
| A-TRR | -0.0947 | -0.4064 | -1.0105 |
| A-$t_{dwell}$ | 0.1207 | 1.6877 | 1.2213 |
| A-TCD | 0.3154 | -1.3361 | -0.0786 |
| A-TCR | -0.0874 | 0.3182 | 0.0236 |
| C-FCR | -0.0419 | -3.2396 | 0.1876 |
| C-TRR | -0.0058 | -2.0477 | 0.2139 |
| C-$t_{dwell}$ | 0.1364 | 9.8689 | -0.6626 |
| C-TCD | 0.3054 | 0.3886 | -0.2051 |
| C-TCR | -0.0932 | 0.1037 | 0.0470 |
| G-FCR | -0.0148 | -1.8844 | -0.5146 |
| G-TRR | 0.0293 | -0.0477 | -0.1174 |
| G-$t_{dwell}$ | -0.0084 | 2.2122 | 0.6320 |
| G-TCD | 0.3606 | -2.5638 | -0.2139 |
| G-TCR | -0.1246 | 0.7211 | 0.0552 |
| T-FCR | -0.0399 | -2.0203 | -0.5606 |
| T-TRR | -0.0572 | 0.2610 | 0.0379 |
| T-$t_{dwell}$ | 0.0975 | 1.1657 | 0.3715 |
| T-TCD | 0.3277 | -1.6544 | -0.0719 |
| T-TCR | -0.1048 | 0.4362 | 0.0563 |

Coefficients for the first three principal components for the 20 parameters (five kinetic properties for each of the four bases) derived from the single-molecule sequencing signal. FCR: full catalytic rate, TRR: tag release rate, $t_{dwell}$: dwell time, TCD: tag capture dwell time, TCR: tag capture rate. Capital letters in front of the kinetic parameter refer to each of the four tagged nucleotides. Each principal component is normalized such that all its coefficients sum up to one.

**Supplementary Table 4b** Principal component analysis coefficients for RPol2.

| Parameters | PC1 | PC2 | PC3 |
|---|---|---|---|
| A-FCR | 0.6024 | 0.1068 | 0.5512 |
| A-TRR | 0.0509 | -0.0457 | -0.2364 |
| A-$t_{dwell}$ | -0.3412 | -0.0401 | -0.1753 |
| A-TCD | -0.7491 | 0.2177 | 0.2159 |
| A-TCR | 0.2020 | -0.0559 | -0.0635 |
| C-FCR | 0.9479 | 0.4592 | -1.0790 |
| C-TRR | -0.3124 | -0.2723 | 0.5496 |
| C-$t_{dwell}$ | -0.3860 | -0.1144 | 0.2470 |
| C-TCD | -0.7603 | 0.2808 | -0.0604 |
| C-TCR | 0.1900 | -0.0562 | -0.0406 |
| G-FCR | 0.8103 | 0.1592 | 0.5834 |
| G-TRR | 0.0137 | 0.0090 | -0.2380 |
| G-$t_{dwell}$ | -0.3934 | -0.0698 | -0.2135 |
| G-TCD | -0.8715 | 0.3057 | 0.3796 |
| G-TCR | 0.2612 | -0.0856 | -0.1105 |
| T-FCR | 0.7807 | 0.1301 | 1.3108 |
| T-TRR | -0.0771 | -0.0892 | -0.5534 |
| T-$t_{dwell}$ | -0.3812 | -0.0210 | -0.3355 |
| T-TCD | -0.8311 | 0.2515 | 0.3783 |
| T-TCR | 0.2443 | -0.0698 | -0.1096 |

Coefficients for the first three principal components for the 20 parameters (five kinetic properties for each of the four bases) derived from the single-molecule sequencing signal. FCR: full catalytic rate, TRR: tag release rate, $t_{dwell}$: dwell time, TCD: tag capture dwell time, TCR: tag capture rate. Capital letters in front of the kinetic parameter refer to each of the four tagged nucleotides. Each principal component is normalized such that all its coefficients sum up to one.

**Supplementary Table 4c** Principal component analysis coefficients for RPol3.

| Parameters | PC1 | PC2 | PC3 |
|---|---|---|---|
| A-FCR | 0.0734 | -0.1693 | 0.1797 |
| A-TRR | 0.0686 | -0.0513 | -0.1464 |
| A-$t_{dwell}$ | -0.0329 | 0.0558 | -0.0157 |
| A-TCD | 0.0447 | 0.5306 | 0.0899 |
| A-TCR | -0.0166 | -0.1461 | -0.0316 |
| C-FCR | 1.2938 | -0.1789 | 0.0391 |
| C-TRR | -0.7098 | -0.1028 | 0.1960 |
| C-$t_{dwell}$ | -0.1564 | 0.0724 | -0.0584 |
| C-TCD | 0.1582 | 0.6036 | 0.0716 |
| C-TCR | -0.0404 | -0.1824 | -0.0343 |
| G-FCR | 0.1109 | -0.1684 | 0.4817 |
| G-TRR | 0.1149 | 0.1058 | -0.2228 |
| G-$t_{dwell}$ | -0.0526 | 0.0215 | -0.0627 |
| G-TCD | 0.0789 | 0.5947 | 0.1862 |
| G-TCR | -0.0227 | -0.1965 | -0.0571 |
| T-FCR | 0.0249 | -0.2918 | 0.5329 |
| T-TRR | 0.0632 | 0.0636 | -0.1716 |
| T-$t_{dwell}$ | -0.0242 | 0.0704 | -0.1021 |
| T-TCD | 0.0329 | 0.5553 | 0.1813 |
| T-TCR | -0.0089 | -0.1861 | -0.0557 |

Coefficients for the first three principal components for the 20 parameters (five kinetic properties for each of the four bases) derived from the single-molecule sequencing signal. FCR: full catalytic rate, TRR: tag release rate, $t_{dwell}$: dwell time, TCD: tag capture dwell time, TCR: tag capture rate. Capital letters in front of the kinetic parameter refer to each of the four tagged nucleotides. Each principal component is normalized such that all its coefficients sum up to one.

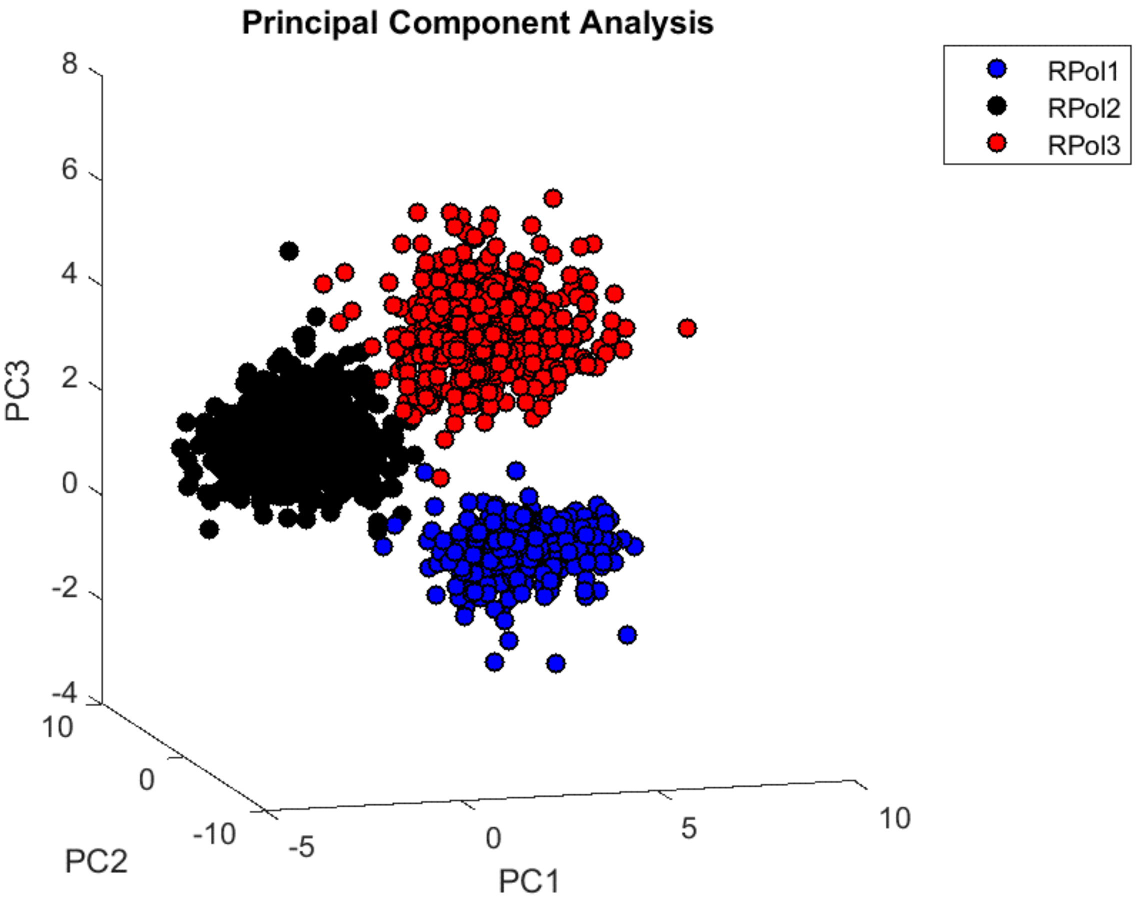


**Supplementary Movie 1** Principal component analysis of polymerase variants. For each polymerase variant (blue: RPol1, black: RPol2, red: RPol3), based on the five derived kinetic properties, this movie shows a well-defined 3D separation between projections of the kinetic parameters onto the first three principal components (PC).

a

**Random 51-mer:**

5'-AGTTCAACCGATCGAGTTCCGACCGGTAAGGTAGCATTGTCTCGCACAATC-3'

**Random 32-mer with flanking 19-nt "universal" primer:**

5'-GGCTAAAATTGTAGAGGTAAATATGTACATATTGCCCGCTCTCCCCACTCT-3'

Color assignment: universal primer barcode

b

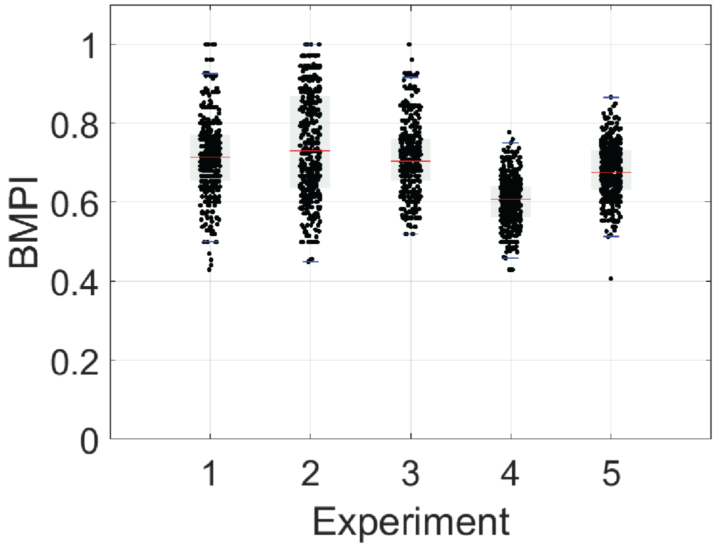


**Supplementary Figure 6** Barcode identification in a pooled sequencing reaction. (**a**) Sequences of the two 51-bp ssDNA templates used as non-matching barcodes. (**b**) Barcode match probability index (BMPI) values in a sequencing experiment, in which each of the three polymerase variants were loaded with a unique barcoded DNA templates (**1**: RPol1-CBT1, **2**: RPol2-CBT2 and **3**: RPol3-CBT3) and pooled in equimolar ratios. (**1-3**) Upon classification, there were more reads with BMPI values >0.80 when compared to the loaded (correct) templates versus the two random templates: (**4**) a random 51-bp barcode or (**5**) a random 32-bp barcode with the universal 19-bp flanking priming site as shown in **a**. This indicates the possibility of barcode identification in a pooled experiment. In each distribution, the red central mark indicates the mean. Raw data are jittered along the x-axis for clarity.

**Supplementary Table 5** Frequency count of barcodes in the pooled, 3-plex sequencing reaction with barcode match probability index (BMPI) value greater than 0.80 as computed by the barcode classifier.

| **BMPI Values** | **RPol1-CBT1** | **RPol2-CBT2** | **RPol3-CBT3** | **Total RPol-CBT** |
|---|---|---|---|---|
| Total Raw Reads | 418 | 390 | 392 | 418 |
| Raw Reads with BMPI > 0.80 | 154 | 66 | 60 | 280 |
| % of Raw Reads with BMPI > 0.80 | 36.8 | 16.9 | 15.3 | 66.9 |

BMPI values were obtained by classifying a 3-plex sequencing experiment data, in which each of the three polymerase variants were loaded with a unique barcoded DNA templates (**1**: RPol1-CBT1, **2**: RPol2-CBT2 and **3**: RPol3-CBT3) and pooled in equimolar ratios.

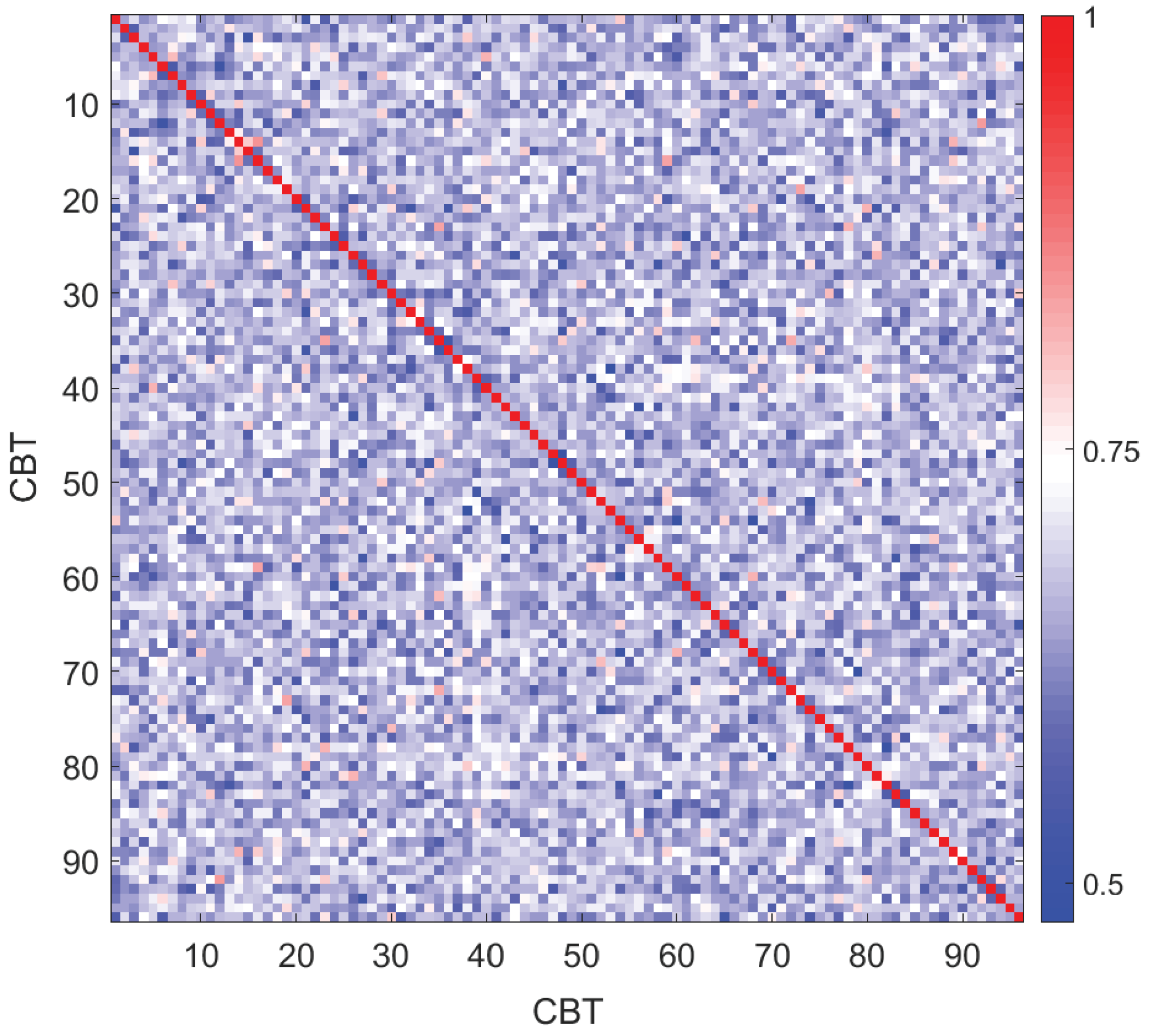


**Supplementary Figure 7** Barcode design for unique identification. Heatmap of sequence identity values of the 96 circular barcoded templates (CBT) calculated by the Smith-Waterman local alignment algorithm. Each barcode sequence (x-axis) was compared to all of the other 96 CBTs in the same barcode set (y-axis) and the sequence identity value was recorded. The probability scale for local alignment is shown on the right, where 0 means total mismatch and 1 denotes total match. The red diagonal line represents perfect identity, when the barcodes are aligned to themselves. For all off-diagonal CBTs, the sequence identities were <85% when the templates were locally aligned to each other.

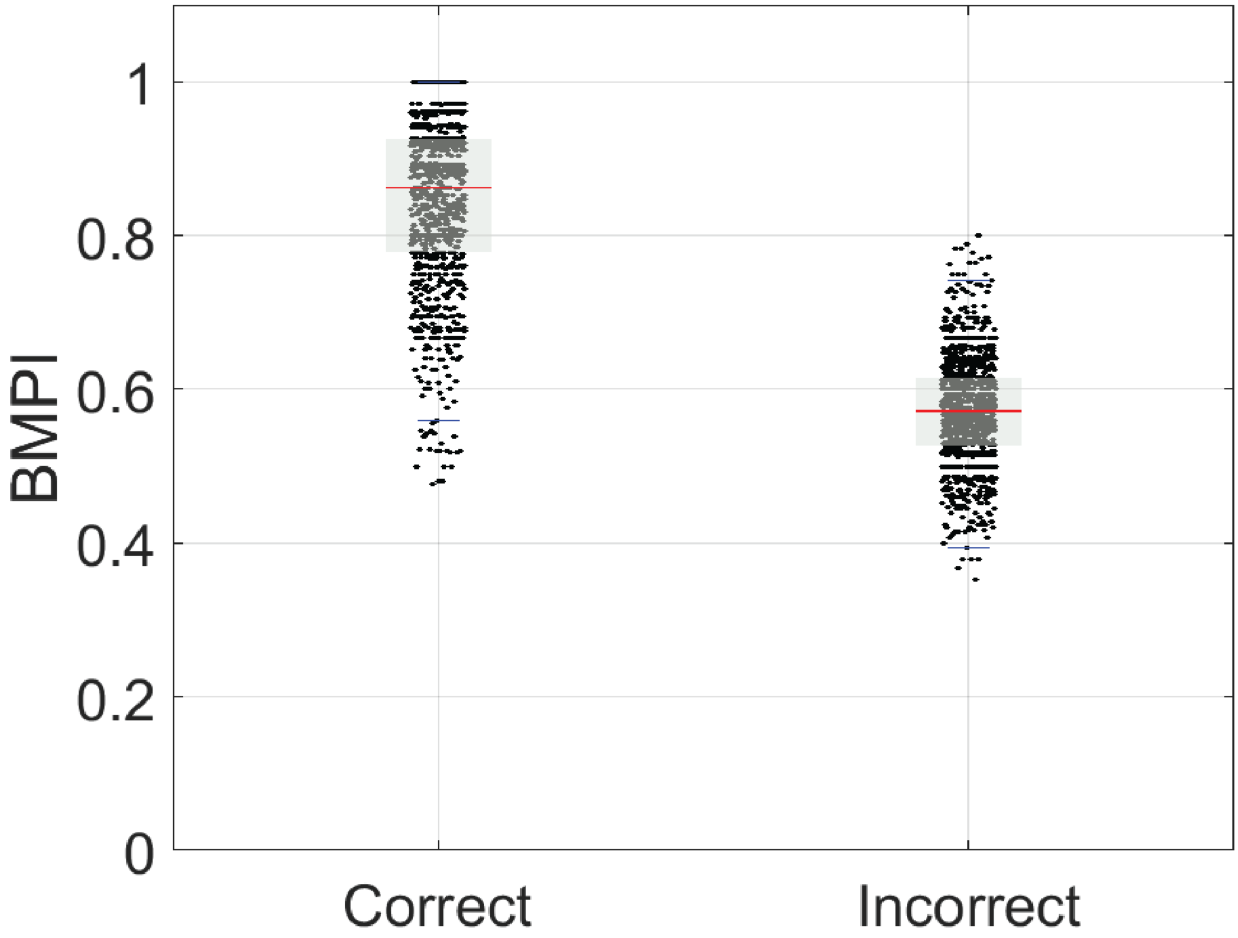


**Supplementary Figure 8** *In silico* demonstration of barcode identification. The left plot shows the calculated BMPI values (black dots) of the randomly sampled quality raw reads from **Fig. 2** compared to the experiment-specific (correct) barcodes. The right plot represents a similar classification of the same raw reads when compared to a randomly chosen barcode (incorrect) out of a list of 96 sequences designed for the high-throughput experiments. Notice the BMPI value decreases from left to right. On each grey box, the red central mark indicates the mean, and the bottom and top edges of the box indicate the 25th and 75th percentiles, respectively. Raw data are jittered along the x-axis for clarity.

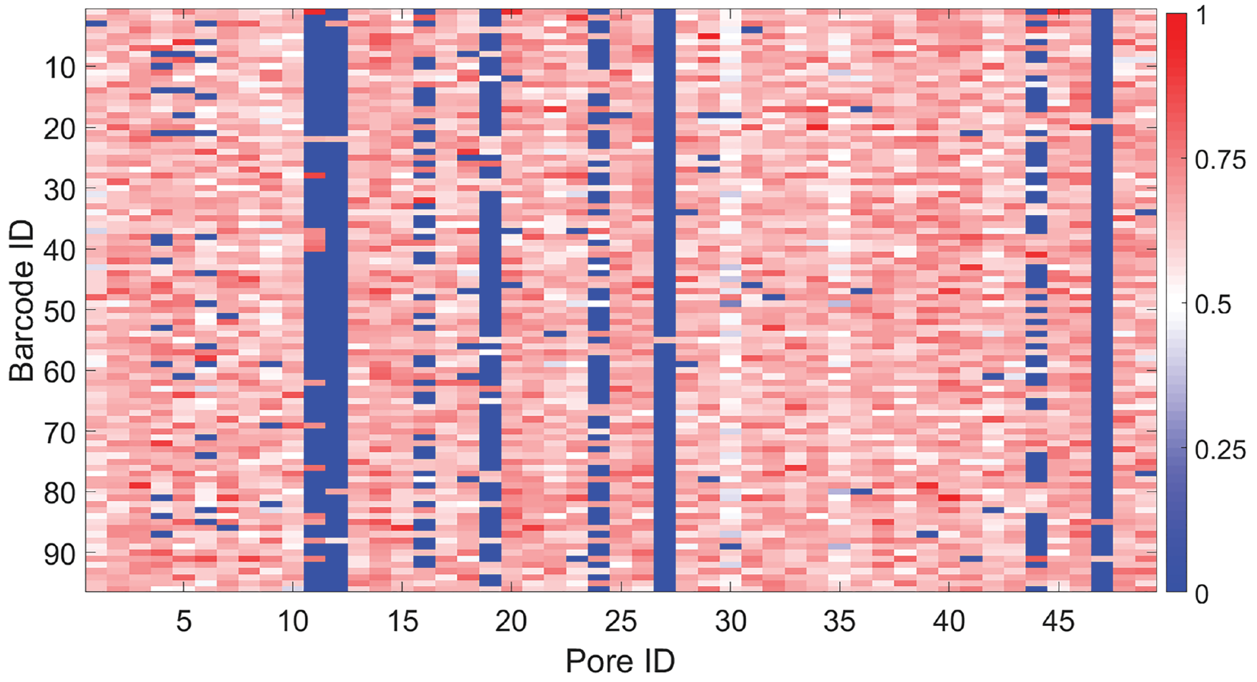


**Supplementary Figure 9** Representative heatmap of raw read classification. Each raw read comes from a single pore-polymerase conjugate complexed with a specific single barcoded template. Each raw read from a pore (x-axis) was compared to all of the 96 CBTs (y-axis) and the barcode match probability index (BMPI) value was recorded (**Methods**). BMPI is a probabilistic measure of barcode identification with a possible range of [0,1], as shown in the scale bar, where 0 means total mismatch and 1 denotes a total match. The maximum scoring BMPI value, above the 0.80 threshold, identified the most likely barcode candidate in each column. A BMPI value of 0 (blue) means that, at the initial classification step, the raw read did not meet the quality read criterion (**Methods**). Reads with maximum BMPI value <0.80 and BMPI values of 0 were discarded from the downstream analysis. Only 50 raw read evaluations are shown here for clarity.

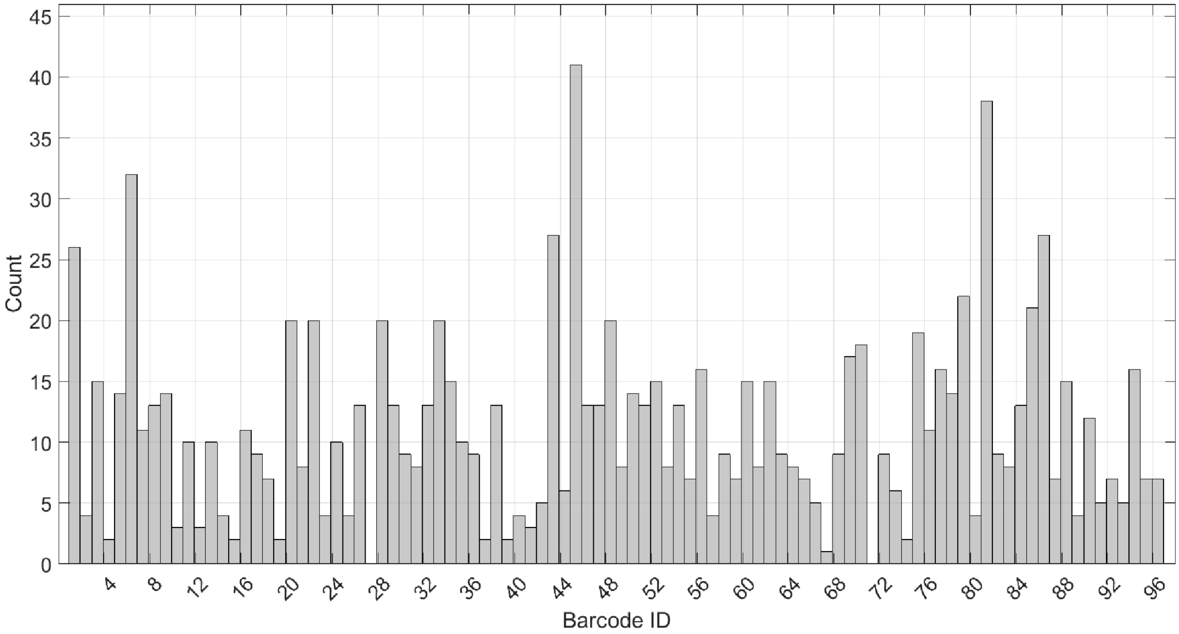


**Supplementary Figure 10** Distribution of experimentally observed barcodes for a single polymerase variant (RPol2). Out of 96 possible barcodes, a total of 94 (98%) were uniquely identified by the alignment-based classification algorithm (**Methods**).

**Supplementary Table 6** Percent probability of misidentifying barcodes in the expected set as another barcode in another set determined by the barcode classifier algorithm.

| **Barcode Counts** | **RPol1:CBT1-32** | **RPol2:CBT33-64** | **RPol3:CBT65-96** |
|---|---|---|---|
| Total Barcodes Identified | 67 | 249 | 383 |
| % of Barcodes in CBT1-32 | *88.06* | 11.65 | 7.83 |
| % of Barcodes in CBT33-64 | 11.94 | *83.94* | 3.39 |
| % of Barcodes in CBT65-96 | 0.00 | 4.42 | *88.77* |
| % of Misidentified Barcodes | **11.94** | **16.06** | **11.23** |

BMPI values were obtained by classifying data from a 96-plex sequencing experiment, in which each of the three polymerase variants were loaded with a unique set of 32 barcoded DNA templates (**1**: RPol1-CBT1-32, **2**: RPol2-CBT33-64 and **3**: RPol3-CBT65-96) and pooled in equimolar ratios. Barcodes, which were sampled at least as many times as the average observation frequency of false positives, were only used in the classification.

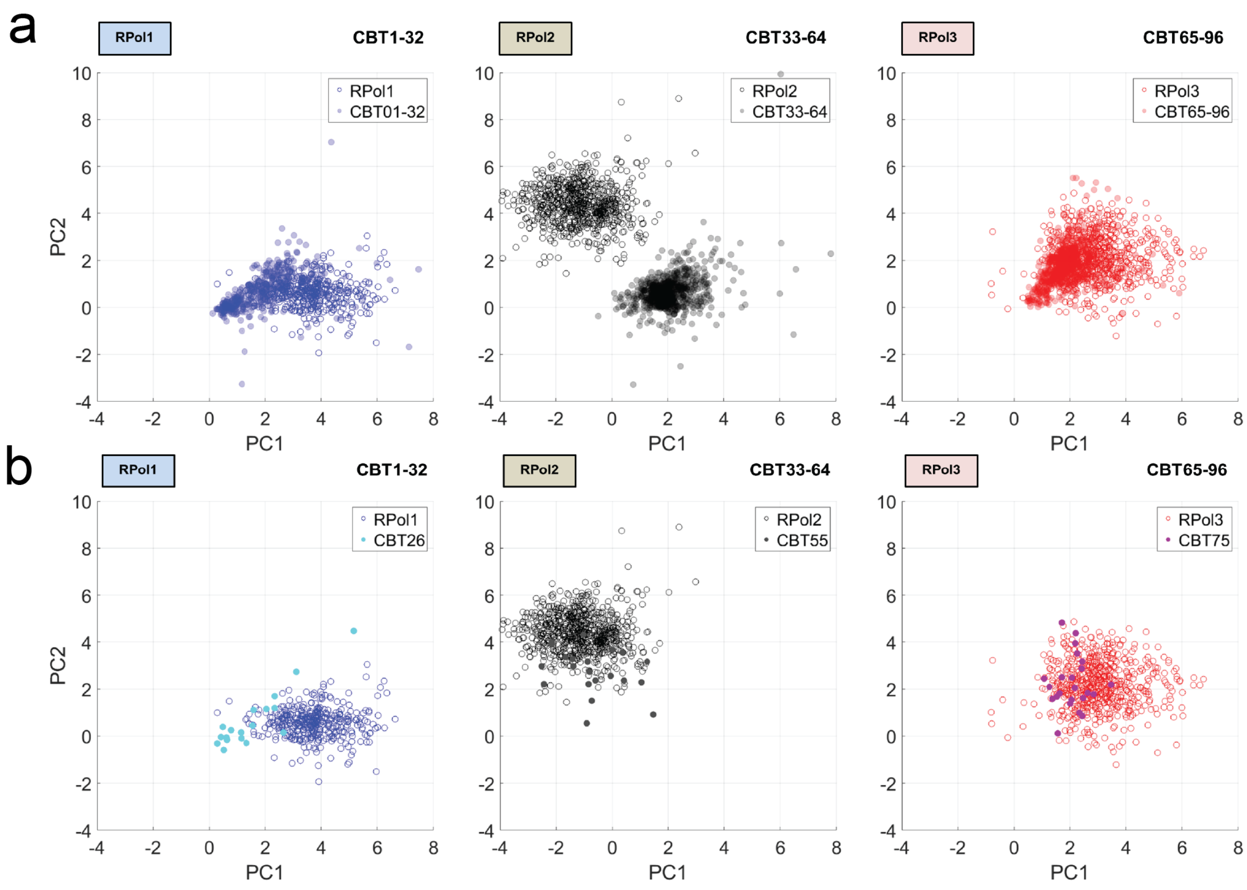


**Supplementary Figure 11** Barcode mapping to kinetic properties. The original PCA clusters, from the singleplex RPol-CBT experiments (**Fig. 5**), are shown as blue, black and red circles, while the barcode mappings in solid circles. (**a**) All barcodes in the first set (CBT1-32), the second set (CBT33-64), and the third set (CBT65-96) are mapped back to the PC1-PC2 plot. (**b**) Representative examples of a single CBT (with high-frequency count) mapping.

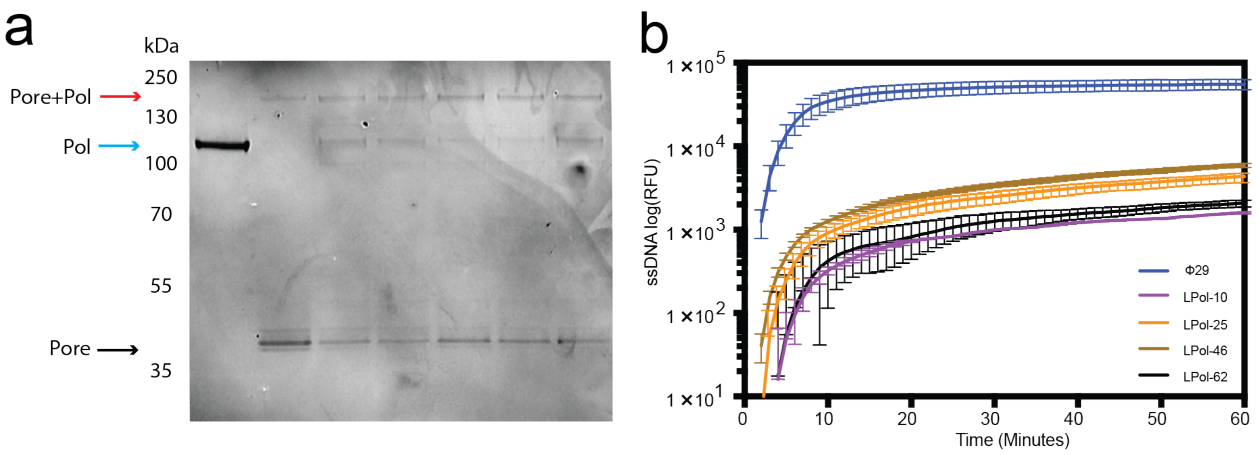


**Supplementary Figure 12** Protein expression and function confirmation. (**a**) 4-12% Bis-Tris protein gel of six randomly picked DNA polymerase (Pol) variants from the library conjugated to αHL pore (Pore). Polymerase alone migrates as a ~100 kDa band (blue arrow). When the polymerase is conjugated with the pore (black arrow), the band shifts to a higher molecular weight indicating that conjugation has occurred (red arrow). (**b**) Rolling circle amplification (RCA) plots of the top four hits of the polymerase variants from the mutagenesis library, indicating that the polymerase activity is maintained after purification as the amplified product RFU follows a logarithmic growth. Positive control (blue curve) contained wild-type ϕ29 DNA polymerase (NEB).

**Supplementary Table 7** Statistical details.

| | μ ± SEM | | | |
|---|---|---|---|---|
| | **LPol-10** | **LPol-25** | **LPol-46** | **LPol-62** |
| **FCR ($s^{-1}$)** | 1.50 ± 0.62 | 2.74 ± 1.09 | 2.20 ± 0.61 | 0.42 ± 0.16 |
| **$t_{dwell}$ (s)** | 0.72 ± 0.17 | 1.79 ± 1.30 | 0.40 ± 0.07 | 1.83 ± 0.61 |
| **TRR ($s^{-1}$)** | 0.67 ± 0.33 | 1.12 ± 0.27 | 1.48 ± 0.53 | 0.63 ± 0.33 |
| **TCR ($s^{-1}$)** | 1.02 ± 0.12 | 1.67 ± 0.54 | 1.45 ± 0.21 | 0.81 ± 0.19 |
| **#** | 3 | 3 | 4 | 3 |

For each of the LPol-CBT top hits, mean (μ) and standard error of the mean (SEM) values corresponding to the full catalytic rate (FCR), dwell time ($t_{dwell}$), tag release rate (TRR) and tag capture rate (TCR) for the nucleotides are shown. #: number of quality reads used in the calculations.

> SpyCatcher-ϕCPV4 DNA Polymerase

MHHHHHHHHSGDYDIPTTENLYFQGAMVDTLSGLSSEQGQSGDMTIEEDSATHIKFSKRDEDGKELAGAT
MELRDSSGKTISTWISDGQVKDFYLYPGKYTFVETAAPDGYEVATAITFTVNEQGQVTVNGKATKGDAHI
GGSDKHTQYVKEHSFNYDEYKKANFDKIECLIFDTESCTNYENDNTGARVYGWGLGVTRNHNMIYGQNLN
QFWEVCQNIFNDWYHDNKHTIKITKTKKGFPKRKYIKFPIAVHNLGWDVEFLKYSLVENGFNYDKGLLKT
VFSKGAPYQTVTDVEEPKTFHIVQNNNIVYGCNVYMDKFFEVENKDGSTTEIGLCLDFFDSYKIITCAES
QFHNYVHDVDPMFYKMGEEYDYDTWRSPTHKQTTLELRYQYNDIYMLREVIEQFYIDGLCGGELPLTGMR
TASSIAFNVLKKMTFGEEKTEEGYINYFELDKKTKFEFLRKRIEMESYTGGYTHANHKAVGKTINKIGCS
LDINSSYPSQMAYKVFPYGKPVRKTWGRKPKTEKNEVYLIEVGFDFVEPKHEEYALDIFKIGAVNSKALS
PITGAVSGQEYFCTNIKDGKAIPVYKELKDTKLTTNYNVVLTSVEYEFWIKHFNFGVFKKDEYDCFEVDN
LEFTGLKIGSILYYKAEKGKFKPYVDHFTKMKVENKKLGNKPLTNQAKLILNGAYGKFGTKQNKEEKDLI
MDKNGLLTFTGSVTEYEGKEFYRPYASFVTAYGRLQLWNAIIYAVGVENFLYCDTDSIYCNREVNSLIED
MNAIGETIDKTILGKWDVEHVFDKFKVLGQKKYMYHDCKEDKTDLKCCGLPSDARKIIIGQGFDEFYLGK
NVEGKKQRKKVIGGCLLLDTLFTIKKIMF

Color assignment: His-Tag SpyCatcher Linker ϕCPV4 Mutation

**Supplementary Figure 13** Sequence of Clostridium phage ϕCPV4 DNA polymerase with colored annotations for the various protein sequence regions. In one of the polymerase mutants identified as our top hits (LPol-46), the lysine highlighted in yellow was mutated to an isoleucine.

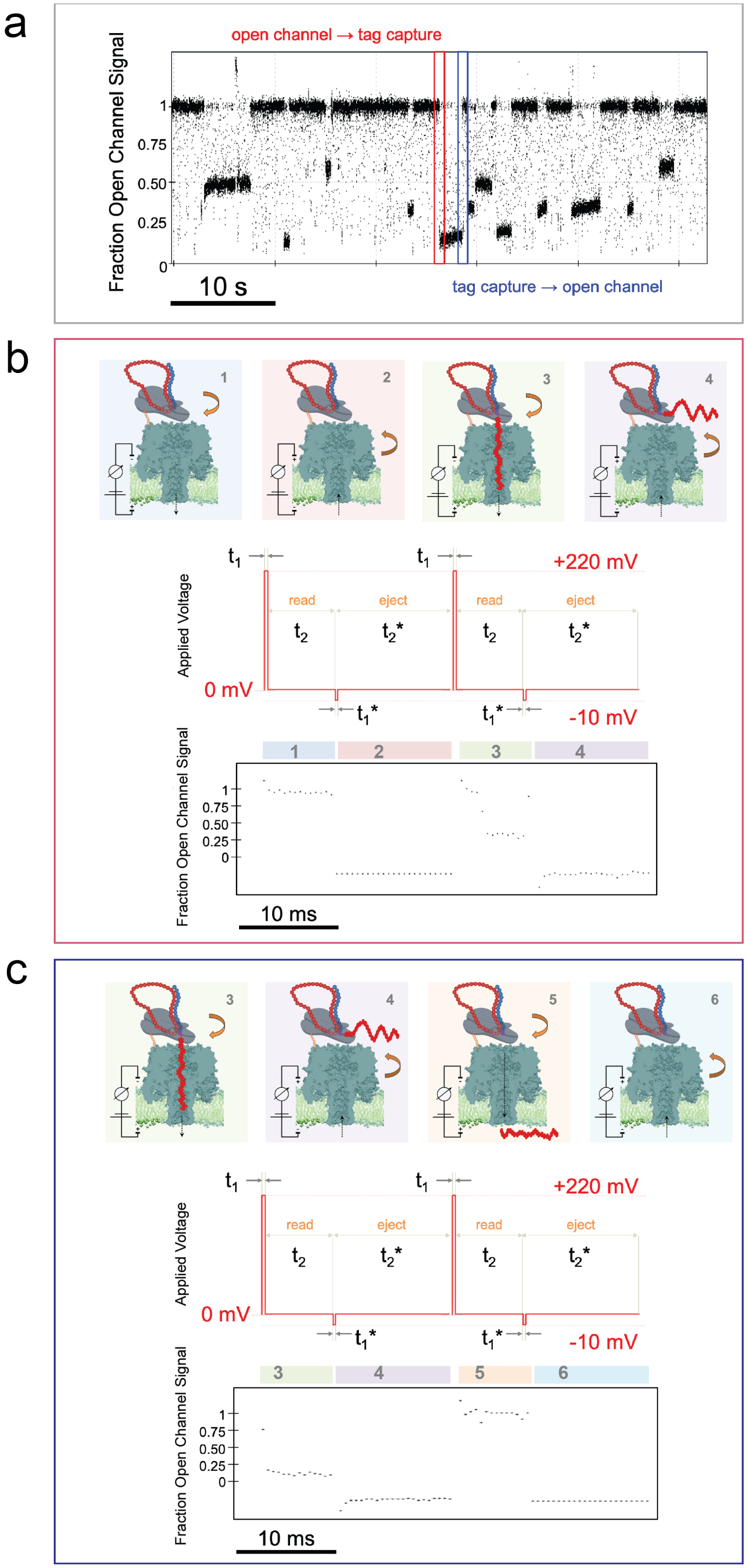
a
Fraction Open Channel Signal
open channel → tag capture
1
0.75
0.50
0.25
0
10 s
tag capture → open channel
b
1
2
3
4
t1
t1
+220 mV
read
eject
read
eject
t2
t2*
t2
t2*
Applied Voltage
0 mV
t1*
t1*
-10 mV
Fraction Open Channel Signal
1
0.75
0.50
0.25
0
10 ms
c
3
4
5
6
t1
t1
+220 mV
read
eject
read
eject
t2
t2*
t2
t2*
Applied Voltage
0 mV
t1*
t1*
-10 mV
Fraction Open Channel Signal
1
0.75
0.50
0.25
0
10 ms

**Supplementary Figure 14** Dynamic voltage control. (**a**) Representative fraction open channel signal (FOCS) *versus* time trace highlighting an open channel to tag capture [red rectangle] and tag capture to open channel [blue rectangle] transitions. (**b**) Voltage commands during open channel to tag capture transition. Positive voltage bias ($V_{min}$=+220 mV) is applied across the nanopore in response to the reset signal. The reset signal is kept high for a short time period ($t_1$=200 μs) during which the capacitor is charged. Then the reset signal is kept low (therefore $V_{min}$=0 mV) for a longer time period ($t_2$=8 ms), so that it is discharged, and the rate of decay is determined. This last voltage command sequence is defined as the "read" period. Immediately after, the same procedure is repeated, but now with a negative voltage bias ($V_{min}$=-10 mV for $t_1$*=200 μs) which will force the captured tag to be ejected from the pore. The subsequent discharge stage ($V_{min}$=0 mV) is defined as the "eject" period with time duration of $t_2$*=12 ms. During a sequencing experiment, the "read" and "eject" commands continuously alternate to repeatedly interrogate the same tagged nucleotide during incorporation as well as determine open channel conditions (inferring the absence of incorporation activity of the polymerase). Schematic of an open channel to tag capture signal transition. **1** represents a "read" period when no tagged nucleotide is present. **2** represents an "eject" period when no tagged nucleotide is present. **3** represents a "read" period when a tagged nucleotide is being incorporated and thus tags are being captured in the pore. **4** represents an "eject" period when a tagged nucleotide is being incorporated and thus tags are being ejected from the pore. The corresponding FOCS response is shown below the control signal. (**c**) Voltage commands during tag capture to open channel transition. The same dynamic voltage control is applied as in **b**. Schematic of a tag capture to open channel signal transition. **3** and **4** as in **b**. **5** represents a "read" period when a tagged nucleotide is being cleaved at the end of the catalytic cycle and thus the tag is being released through the pore. **6** represents an "eject" period when a tagged nucleotide has been released thus not present in the pore. The corresponding FOCS response is shown below the control signal.

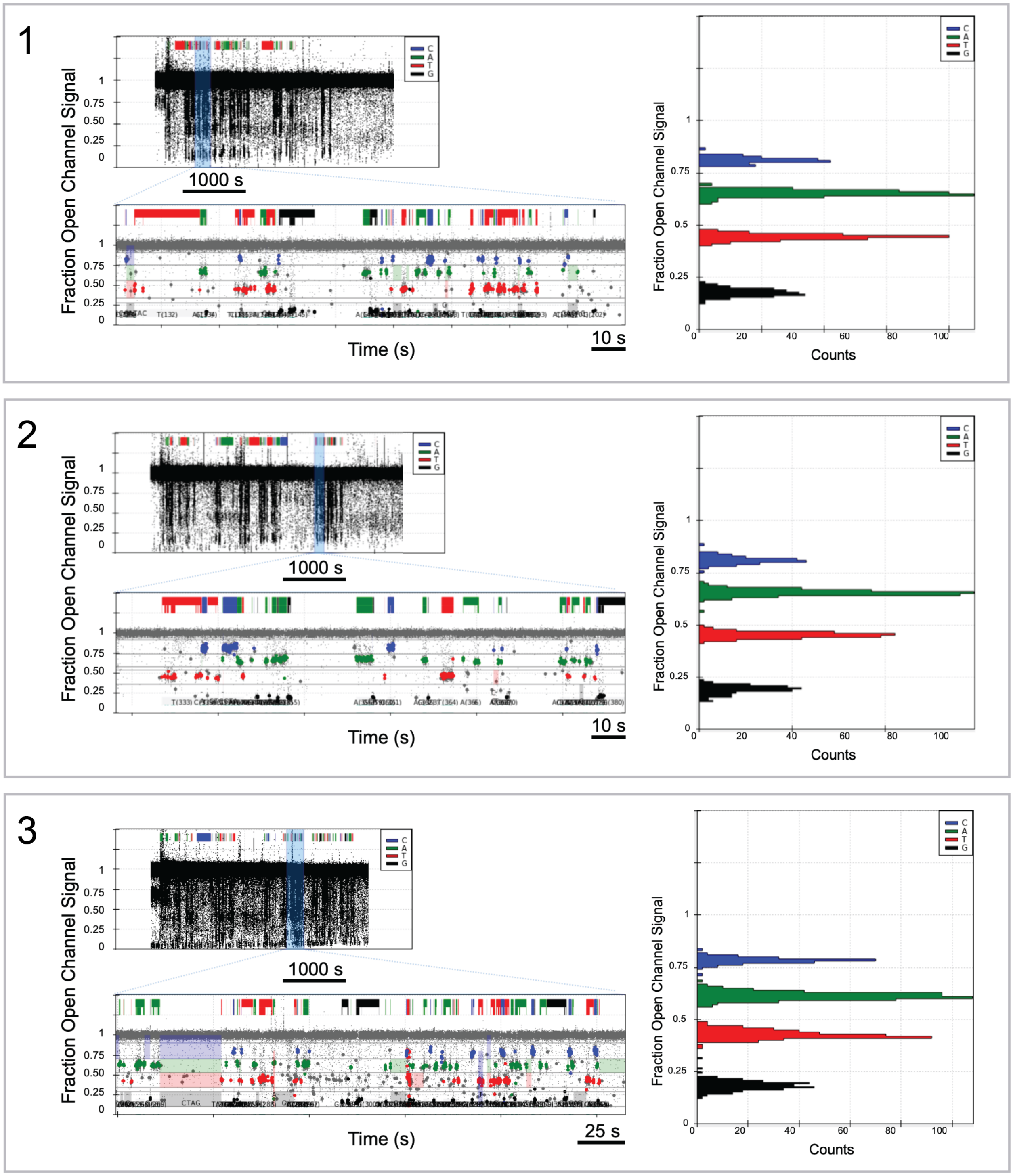
1
Fraction Open Channel Signal
1
0.75
0.50
0.25
0
1000 s
C
A
T
G
Time (s)
10 s
Counts
0
20
40
60
80
100
2
Fraction Open Channel Signal
1000 s
Time (s)
10 s
Counts
3
Fraction Open Channel Signal
1000 s
Time (s)
25 s
Counts

**Supplementary Figure 15** Typical fraction open channel signal (FOCS) traces of multiple pores (**1**, **2**, and **3**). FOCS *versus* time trace of tagged nucleotide captures for a single pore during a typical DNA sequencing experiment [left, top panels]. Base calls are highlighted in standard Sanger colors [left, bottom panels]. Histogram of the FOCS is shown in the right sub-panels. The four peaks represent signal associated with each tagged nucleotide capture.

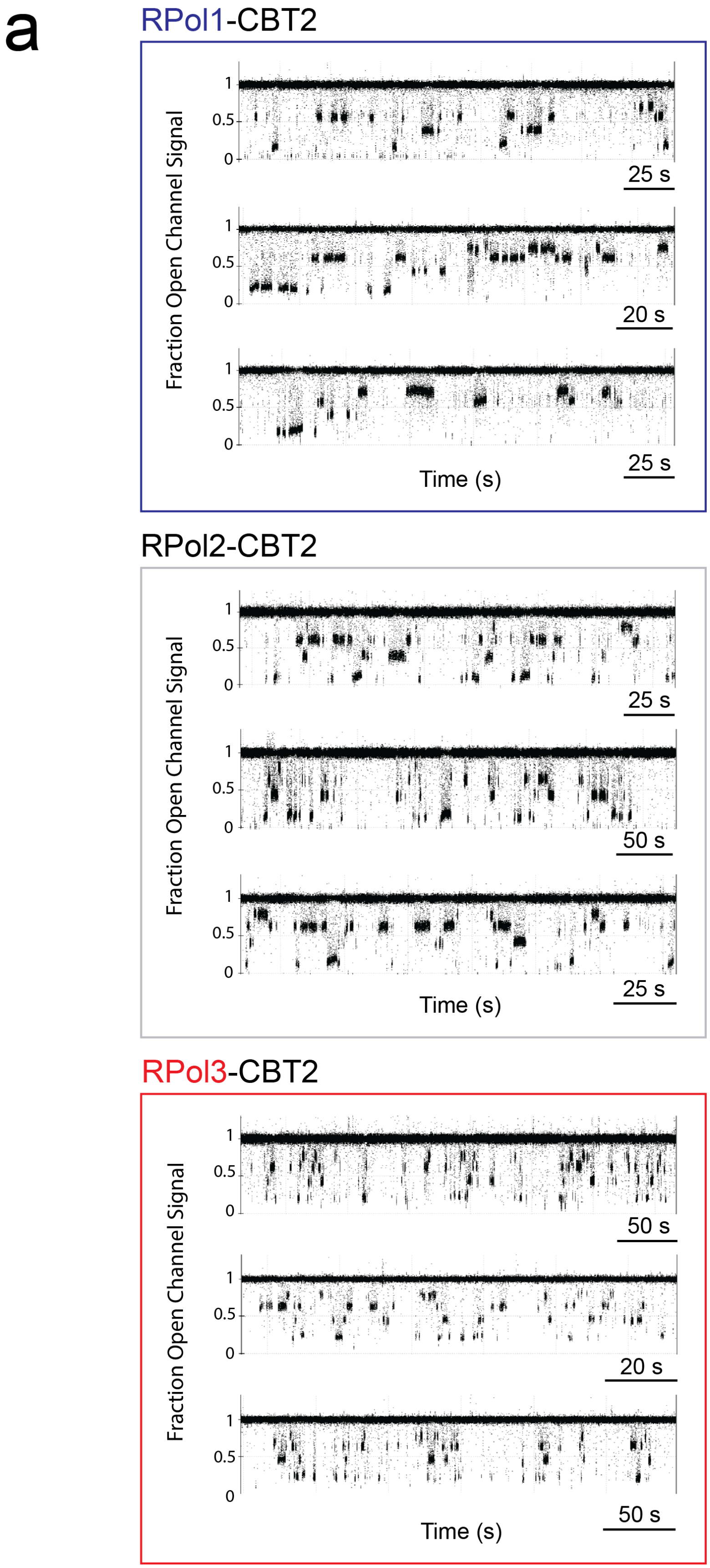
a
RPol1-CBT2
Fraction Open Channel Signal
1
0.5
0
25 s
20 s
25 s
Time (s)
RPol2-CBT2
Fraction Open Channel Signal
25 s
50 s
25 s
Time (s)
RPol3-CBT2
Fraction Open Channel Signal
50 s
20 s
50 s
Time (s)

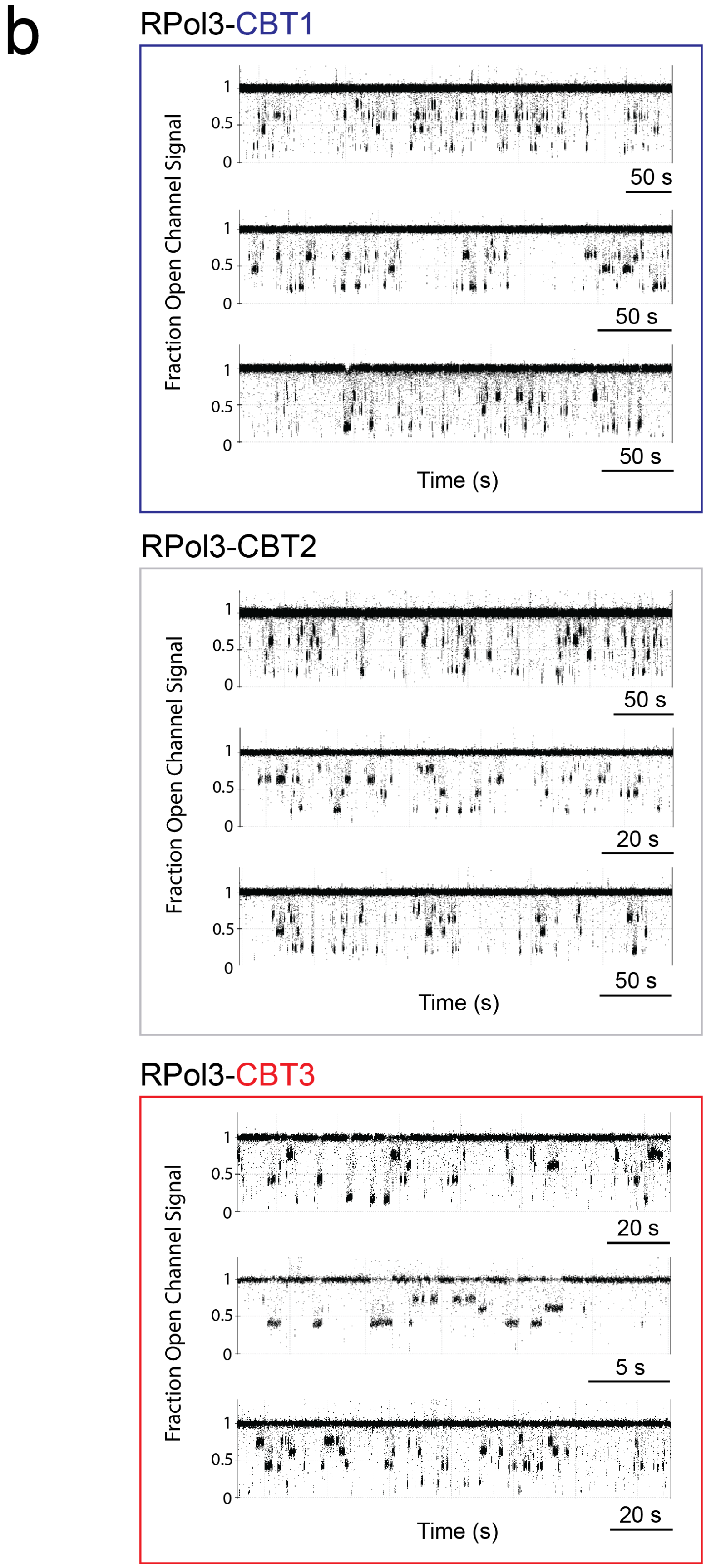
b
RPol3-CBT1
Fraction Open Channel Signal
1
0.5
0
50 s
50 s
50 s
Time (s)
RPol3-CBT2
Fraction Open Channel Signal
50 s
20 s
50 s
Time (s)
RPol3-CBT3
Fraction Open Channel Signal
20 s
5 s
20 s
Time (s)

**Supplementary Figure 16** Example traces of experimental repeats. (**a**) Fraction open channel signal (FOCS) trace examples comparing different polymerases (RPol1, RPol2 and RPol3) loaded with the same template (CBT2). (**b**) FOCS trace examples directly comparing the same polymerase (RPol3) with different templates (CBT1, CBT2 and CBT3). For each RPol:CBT assignment the data was obtained from individual pores of different measurements.
